# Integration of Multi-level Molecular Scoring for the Interpretation of RAS-Family Genetic Variation

**DOI:** 10.1101/794131

**Authors:** Swarnendu Tripathi, Nikita R. Dsouza, Raul A. Urrutia, Michael T. Zimmermann

## Abstract

Protein-coding genetic variants are the first considered in most studies and Precision Medicine workflows, but their interpretation is primarily driven by DNA sequence-based analytical tools and annotations. Thus, more specific and mechanistic interpretations should be attainable by integrating DNA-based scores with scores from the protein 3D structure. However, reliable and reproducible standardization of methods that use 3D structure for genomic variation is still lacking. Further, we believe that the current paradigm of aiming to directly predict the pathogenicity of variants skips the critical step of inferring, with precision, molecular mechanisms of dysfunction. Thus, we report herein the development and evaluation of single and composite 3D structure-based scores and their integration with protein and DNA sequence-based scores to better understand not only if a genomic variant alters a protein, but how. We believe this is a critical step for understanding mechanistic changes due to genomic variants, designing functional validation tests, and for improving disease classifications. We applied this approach to the RAS gene family encoding seven distinct proteins and their 935 unique missense variants present somatically in cancer, in rare diseases (termed RASopathies), and in the currently healthy adult population. This knowledge shows that protein structure-based scores are distinct from information available from genomic annotation, that they are useful for interpreting genomic variants, and they should be taken into consideration in future guidelines for genomic data interpretation.

**Significance Statement:** Genetic information from patients is a powerful data type for understanding individual differences in disease risk and treatment, but most of the genetic variation we observe has no mechanistic interpretation. This lack of interpretation limits the use of genomics data in clinical care. Standard methods for genomics data interpretation take advantage of annotations available for the human reference genome, but they do not consider the 3D protein molecule. We believe that changes to the 3D molecule must be considered, to augment current practice and lead to more precise interpretation. In this work, we present our initial process for systematic multi-level molecular scores, including 3D, to interrogate 935 RAS-family variants that are relevant in both cancer and rare diseases.

## Introduction

To interpret genomic variants, currently used clinical genomics guidelines rely on recurrent observations in diseases or tumors compared to the general healthy population, and inferred impact on the encoded protein [1, 2]. However, in our view, the gene product itself must take center stage of the analytical process, either using bioinformatics, functional validation, or both. The fundamental concept underlying this idea is that the structure and dynamics of RNA and proteins ultimately determine whether a variation can be tolerated or becomes pathogenic. Further, current approaches aim to directly predict pathogenicity resulting in different levels of predictive performance [3-5]. We believe that this approach bypasses the necessary step of determining the molecular mechanism of dysfunction. Thus, the wide adoption of protein 3D structure and time-dependent dynamics (4D) is of paramount importance since the mechanistic interpretation of novel genetic variants will ultimately be inferred from the study of the gene product.

Genetic variants activating Rat sarcoma (RAS) genes are among the most recurrent somatic alterations in human cancers, affecting up to 25% of solid tumors [6]. The RAS family of small GTPases has 31 members and all act as signal transducers influencing cellular growth and differentiation. Genetic variants in RAS proteins, when present in the germline, are responsible for rare congenital diseases known as RASopathies [7, 8], such as Noonan (RRAS) and Costello (HRAS) Syndromes. The most clinically relevant, due to their common oncogenic mutation, are KRAS, HRAS, and NRAS [9], discovered from the study of oncogenic viruses and neuroblastomas [10]. Additional family members have been identified through sequencing of tumors (MRAS), RASopathy patients, and neural transformation (e.g. RERG and RRAS2). In cancer, there are two highly recurrent RAS activating variant sites, referred to as hotspots, but many genetic alterations are observed outside of the hotspot sites somatically in cancer, in rare diseases, and in the currently healthy adult population. This genetic spectrum differs for each member of the RAS family. Experimental data indicates that each type of alteration at the hotspots can lead to different downstream effects [11-13]. Further, non-hotspot variants may not alter the protein in the same way as hotspot mutations and therefore may not have the same implications for clinical management. Thus, better methods to evaluate how genetic variation affects RAS, within and outside of the hotspots, are needed in order to interpret their functional impact.

Recurrent cancer variants such as KRAS G12D have been extensively studied by laboratory experimental and computational methods [14]. However, the full spectrum of rare disease and cancer variants has not been evaluated with the same rigor. In the past, genomic information was less abundant, leading to a bias in the field to only focused on the characterization of variants which define hotspots. However, with the NexGen revolution and its application to medicine, genetic variation outside of the hotspots is observed frequently and understanding their health implications requires further investigations. Therefore, the goal of this study is to help fill the gap in knowledge by assembling and testing a more comprehensive series of computational 3D scores integrated with DNA and protein sequence-based scores (**Figure 1**), for interpreting and classifying the most likely underlying mechanisms of alteration by genetic variants in members of the RAS family of proteins. Our results show that no single score is likely to capture enough detail to fully interpret the effects of RAS genomic variants. Rather, different scores indicate alterations of specific properties or functional roles and their combination, in a distinct manner, is more informative for identifying which properties are altered by the broad spectrum of genetic variation in RAS.

**Figure 1:**
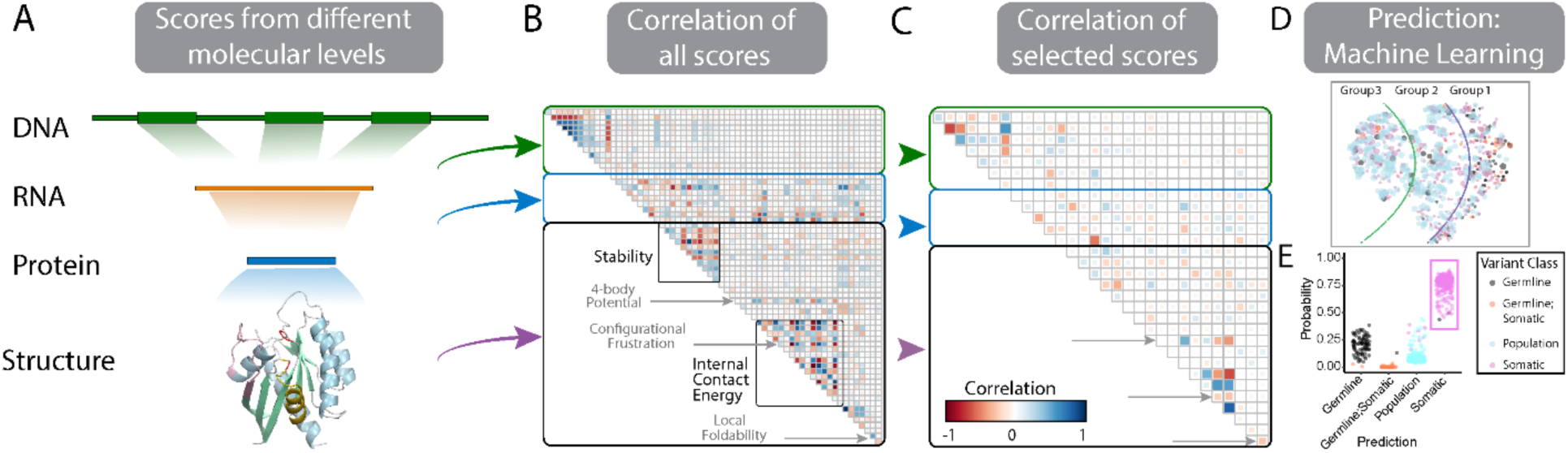
Integrating diverse scores from multiple molecular levels to interpret the effects of genomic variants. **A)** There are at least four distinct molecules that carry relevant information for directly interpreting the effects of genomic variants: the DNA itself, the encoded mRNA, the linear protein, and the 3D folded protein. **B)** Existing clinical solutions to genomic variant interpretation rely on genomic annotations – on observations like recurrence in affected cases, inheritance patterns, and scores derived from mining genetic associations. We have assembled many scores for how genomic variants may alter sequence or structure and analyzed their interrelationships. Larger and labeled versions of the correlation matrices are available in **Figures S4**. We highlight two sections of structure-based scores that have nearly no overlap with one another or with information available in DNA annotations. **C)** Using correlation patterns among scores, reduced the number of individual scores to the 31 that are most unique and therefore most efficiently cover the broadest diversity of properties. The locations of same three scores specifically named in (**B**) are indicated by arrows. **D)** We used the many individual scores to assess the differences among RAS variants and identified groups of variants that affect the protein in similar ways. We believe each of these groups in the reduced dimensional space will represent a different mechanism of dysfunction. **E)** Using the same data, we trained a machine learning classifier to distinguish from among germline pathogenic (RASopathies), somatic, and non-disease genetic variants, to learn which patterns of altered features best associated with pathogenicity.

## Results

### Defining the 3D Landscape of Genomic Variation in Ras Family Members

For the current study, we identified the coding regions in the human genome for the GTPase domains of seven RAS family proteins for which we have 3D protein models (**Table S1 and Figure S1**). We used bioinformatic tools and large publicly available databases to gather all observed genomic variants within these regions and annotated their missense effects (see **Methods** for detailed description). In total, we identified 1103 unique protein-coding genomic variants. Of those variants, 136 affected degenerate codons, leading to our final dataset of 935 unique missense variants from across the seven RAS proteins (**Supplemental data file**). Using this dataset, we sought to contribute to the standardization of structure-based scores by first considering parameters that reflect the landscape of genetic variation in the RAS family proteins, and their impact in the field of Clinical Genomics, Cancer Precision Medicine, as well as hypothesis-based guidance for functional validation. The GTPase domain of RAS family proteins is well studied and contains two conformational switch regions that are critical indicators of activity [8] (**Figure 2A**). Somatic hotspot residues affect these conformational switches (**Figure 2B**). RAS is part of a signaling cascade that results in the phosphorylation of proteins, which in turn also physically interact with RAS, often involving one of the switch regions. Because the physical interaction between proteins is a necessary part of RAS function, we gathered experimental structures of RAS bound to four classes of proteins and identified shared and protein-specific protein-protein interaction surfaces (**Figure 2C and S2**). This knowledge informed the design of our study that combines the analysis of structural properties of regulatory regions, the impact of well-characterized somatic activating mutations, and protein binding interfaces, to characterize which combination of structure-based scores better annotate the likely function of each unique variant.

**Figure 2:**
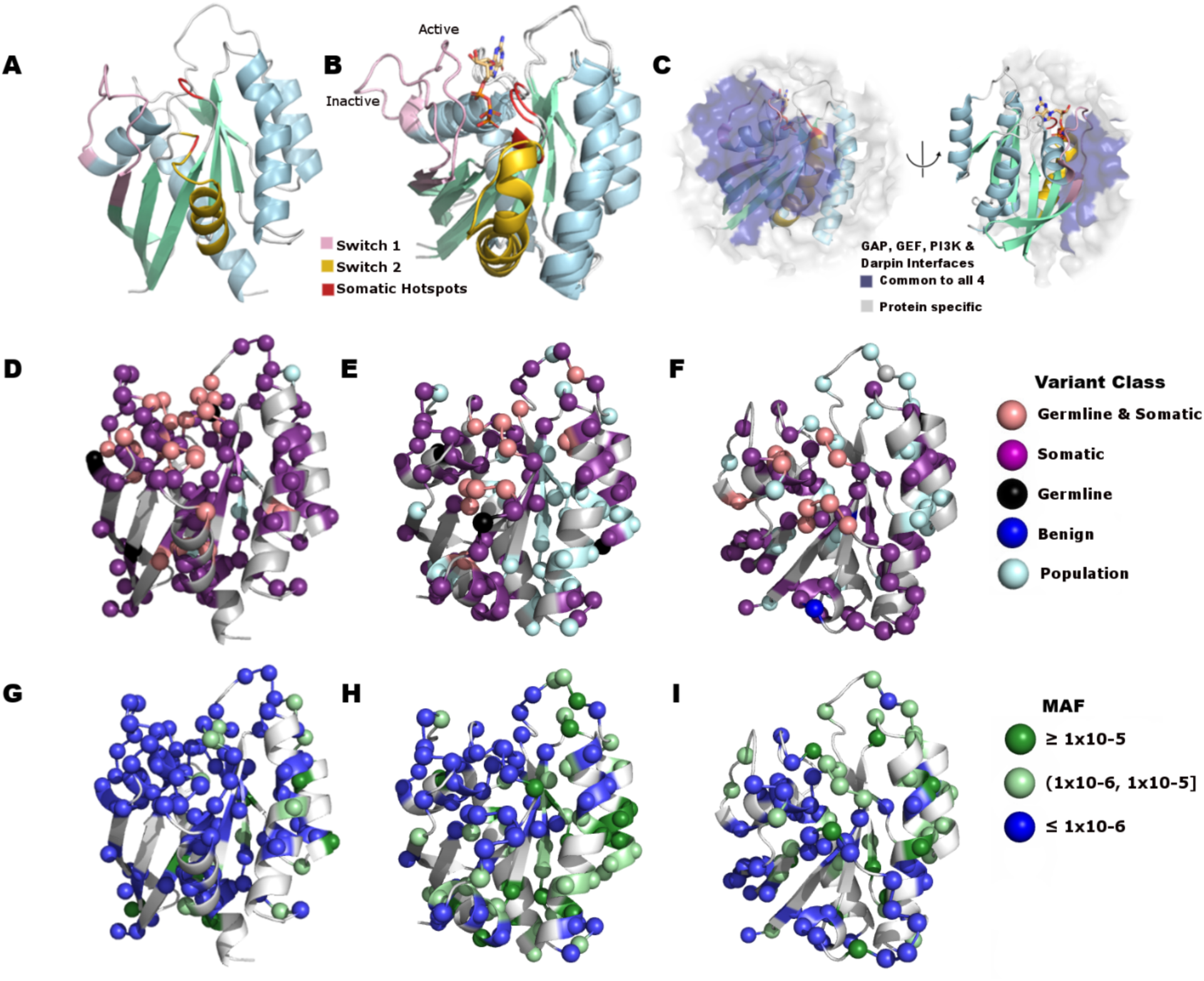
Congenital and Cancer genetic variants have a different profile across RAS proteins. We show three of the seven RAS proteins used in this study. **A)** The structure of KRAS is shown, colored by secondary structure and functional region. **B)** GTP-bound and GDP-bound HRAS differ in conformations of Switch-I and Switch-II. These conformational changes are hallmarks of RAS activity. **C)** RAS proteins function within multi-protein complexes. Protein-complex relationships determined from experimental structure determination are shown as transparent surfaces; those common to four interaction partners are colored blue those that are protein-specific are colored white. Images for each protein interaction are shown in **Figure S2**. **D-I)** Somatic hotspot sites (G12-G13 and Q61) are frequently altered across human solid tumors, but many amino acids throughout the protein are altered somatically. Germline variants responsible for congenital diseases occur in a focal region of the protein structure. This 3D pattern is shared by the RAS family. We show (**D, G**) KRAS, (**E, H**) HRAS, and (**F, I**) NRAS, where amino acids are colored according to their **D-F**) variant class and (**G-I**) MAF in the currently healthy population.

We subsequently mapped these variants to structural models of 7 RAS proteins and identified 935 unique protein variants (**Figure 2 D-I and S3**). Our structural scoring of this large number of variants revealed that while there exist three somatic hotspot residues (G12, G13 and Q61), the genetic diversity in RAS that remains to be evaluated is substantially broader. In addition, 3D patterns of variation show similarities across the family, consistent with conserved functional features, but also has protein-specific patterns (**Figure 2 D-I and S3**). Therefore, we further analyzed the protein interaction surfaces (**Table S2**) and demonstrated that KRAS shows the fewest genetic variants, across germline samples, at protein interaction surfaces (8% of observed variants), with HRAS and NRAS showing few (25%) and other RAS-family proteins showing more (45-85%). Further, variants exclusively observed in the currently healthy adult population very rarely occur at KRAS protein interaction surfaces, rarely for HRAS and NRAS, but not uncommonly for MRAS, RRAS, RRAS2, and RERG. Genetic variants known to cause Germline diseases are nearly non-existent (0-3%) at the protein interaction surfaces of MRAS, RRAS, RRAS2 and RERG, while many occur in KRAS, HRAS, and NRAS (13-24%). Finally, observing the 3D locations of genetic variants, many fall in the switch regions (Switch I: 72 variants; Switch II: 107 variants), cancer hotspots (72 variants), and protein interfaces (465 variants). Because part of switch regions and hotspot sites overlaps with the protein interfaces, a total of 491 unique variants occur across these sites. This result emphasized the need for a more comprehensive approach to scoring because even if we assumed that all variants in each region of the protein had the same effect, though likely they do not, 47.5% would remain uninterpreted. Therefore, these analyses defined the translated landscape of genomic variation in RAS proteins, which should be carefully analyzed and compared in a parametric manner in order to standardize scores for their interpretation.

### Evaluating Relationships Among Sequence- and Structure-Based Scores

We evaluated the single and combinatorial performance of distinct scores that account for data from the DNA sequence, protein sequence, and 3D protein structural levels, which can be used for defining the landscape of human variation (**Figure 1**). We performed a comparative analysis of both sequence-derived and structure-based information for all 935 observed missense genetic changes (**Figure 1A**). At each level, we chose a subset of scores that each provide information about a different type of property (**Figure 1B and S4A**). Detailed descriptions of the scores are provided in the **Methods** section. We selected 31 scores based on the correlation structure between the scores (when individually applied to all variants), and their assessment of different biophysical or biochemical properties, so that they are unique and most efficiently cover the broadest diversity of RAS properties (**Figure 1C and S4B**). Among the 31 are seven DNA sequence-based scores, five protein sequence-based, and 19 protein structure-based scores. Among the 19 structure-based scores, we quantified solvent accessible surface area, global folding stability (and related components), four-body contact potential (to account three dimensional interactions associated with residue packing), local energetic frustration (to identify potential changes in local stability), and local foldability (to define thermodynamic subdomains). Since clinical genomics workflows use annotations and computational (a.k.a. *in silico*) predictions based on DNA information, the information made available by molecular modeling methods, when made stringently parametric and standardized, will bring new opportunities for both diagnosis and experimentation.

Using these approaches, we found a strong correlation among the DNA sequence-based scores (median absolute Spearman correlation coefficient ± median absolute difference = 0.24 ± 0.23), weak correlation between them and protein sequence-based scores (0.07 ± 0.05), and almost no correlation between them and protein structure-based scores (0.04 ± 0.04; **Figure 1B and S4A**). Protein sequence-based scores had moderate correlation with structure-based scores (0.12 ± 0.10). Together, this data demonstrate that parameters inferred from 3D structures of RAS are distinct from those derived from both types of sequence analyses. Among structure-based scores, some have weak or no correlation with any other score, including other 3D scores, indicating their unique assessment of a protein property. For example, differences in ‘ionisation energy’ do not correlate with any other scores (0.04 ± 0.03), nor do local ‘Residue density’ (0.03 ± 0.02) or and ‘Cooperativity Ratio’ (CR) (residue: 0.05 ± 0.04; side-chain: 0.04 ± 0.04). Further, scores quantifying folding stability and energetic frustration provide complementary information to one another (0.21 ± 0.24 and 0.24 ± 0.25, respectively). Thus, we conclude that sequence-based information is insufficient for interpreting genomic variants making the current widespread use of each individually, insufficient. It is imperative that 3D scores be parameterized and integrated with sequence-based scores, to improve genomic data interpretation.

### Individual Scores Reveal Partial Correlations but Do Not Classify RAS Variants

We analyzed patterns among individual scores according to the variant classes of ‘Germline’, ‘Population’, ‘Somatic’, ‘Germline & Somatic’, and VUS (see **Methods** for detailed definitions). We illustrate the distribution of six scores based on sequence (**Figure 3A-B**), amino-acid properties (**Figure 3C-D**) and 3D protein structure (**Figure 3E-F**) and their relative value by variant class. Distributions of other 25 scores by variant class are shown in **Figure S5**. Interestingly, the evolutionary trace (ET) score revealed that the variant sites from Germline & Somatic class are highly conserved (**Figure 3A**) at the protein sequence level. Some degree of conservation is also evident from Germline variant sites. For CADD, a higher average score (more likely altering protein function) is observed for all three disease-associated variant classes (Germline median ± MAD: 3.85 ± 0.75, Somatic: 3.82 ± 0.73, and Germline & Somatic: 3.82 ± 0.53), compared to VUS (3.64 ± 0.78) and Population (3.55 ± 0.93). We observed an increase in heat capacity particularly for the VUS, among germline variants known to cause diseases (Germline), and among both germline and somatic variants that cause diseases (Germline & Somatic). Distributions of the difference in electron-ion interaction potential is more subtle and show opposite behavior for the Germline versus Germline & Somatic and VUS variants. Structure-based scores such as heat capacity, electron-ion interaction potential, hydrophobic solvation free energy, and four-body potential show distinct patterns among variant classes (**Figure 3B-F**). Thus, certain individual scores from each molecular level have associations by variant class, but none clearly classify RAS variants.

**Figure 3:**
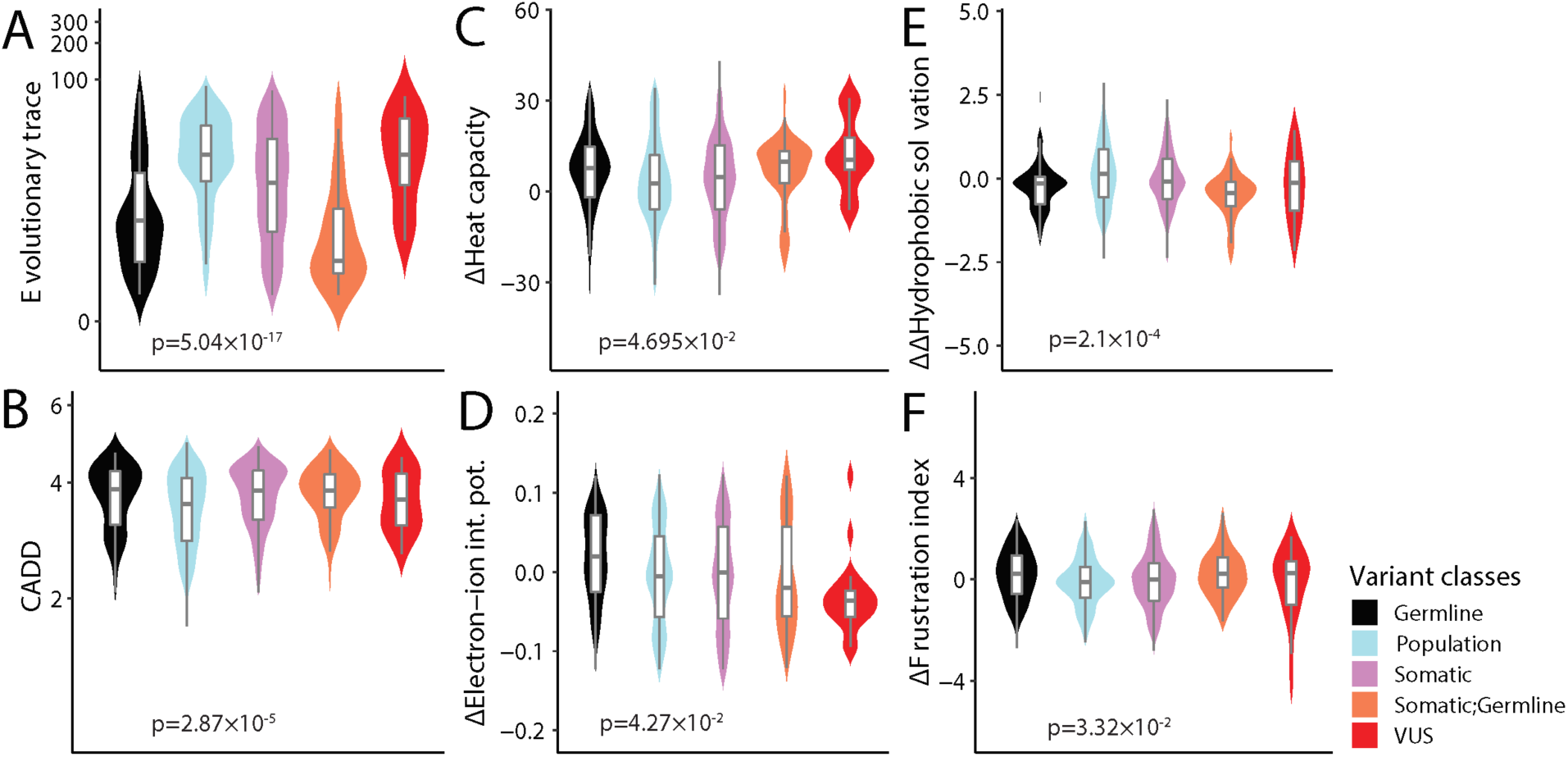
Individual scores do not significantly classify the variants. **A-F)** Distributions of selected scores at different molecular levels are shown in violin plots for five variant classes indicated by different colors. (**A, B**) Sequence-based scores, (**C, D**) difference in the amino-acid properties-based scores and (**E, F**) difference in the protein 3D structure-based scores. The grey horizontal lines, boxes and vertical lines represent the medians, interquartile ranges and 95% confidence intervals of the scores’ distributions (for comparing medians), respectively. For the evolutionary trace score (ET) lower value indicates higher degree of conservation of the sites and for CADD (raw score) the larger the score the more likely the variant has damaging effect. For the amino-acid properties-based and protein structure-based scores, differences were calculated between the variant and the WT. One-way analysis of variance (ANOVA) p-value by variant class is indicated inside each plot. Plots of other 25 scores are shown in **Figure S5**.

Subsequently, we examined the patterns between pairs of scores (**Figure 4**) to define whether and how they distinguish among variant classes. We note that the Population variants with minor allele frequency (MAF) < 1×10^−4^ and ET score < 25 that overlap with Germline, Somatic, and Germline & Somatic variants are not only highly conserved and yet predicted to be altering protein function by the SIFT score (**Figure 4A**). Differences in protein sequence-based heat capacity were negatively correlated with the difference in structure-based hydrophobic solvation free energy (**Figure 4B**). This finding shows that increased (decreased) heat capacity of a variant associates with favorable (unfavorable) changes in hydrophobic solvation free energy. Changes in the structure-based frustration index are negatively correlated with the structure-based four-body potential per residue (**Figure 4C**) suggesting interplay between global and local change in energetics of the variants. Combined, these analyses reveal, 1. correlations among individual scores that reflect molecular properties predicted by knowledge-based and mathematical forcefields, 2. a higher level of predictive knowledge on molecular mechanisms could be gained from their integration, and 3. none of these scores, individually, aids in developing a congruent classification of genomic variants. Therefore, this information is of significantly importance for the development and or optimization of methods that advance genomic data interpretation.

**Figure 4:**
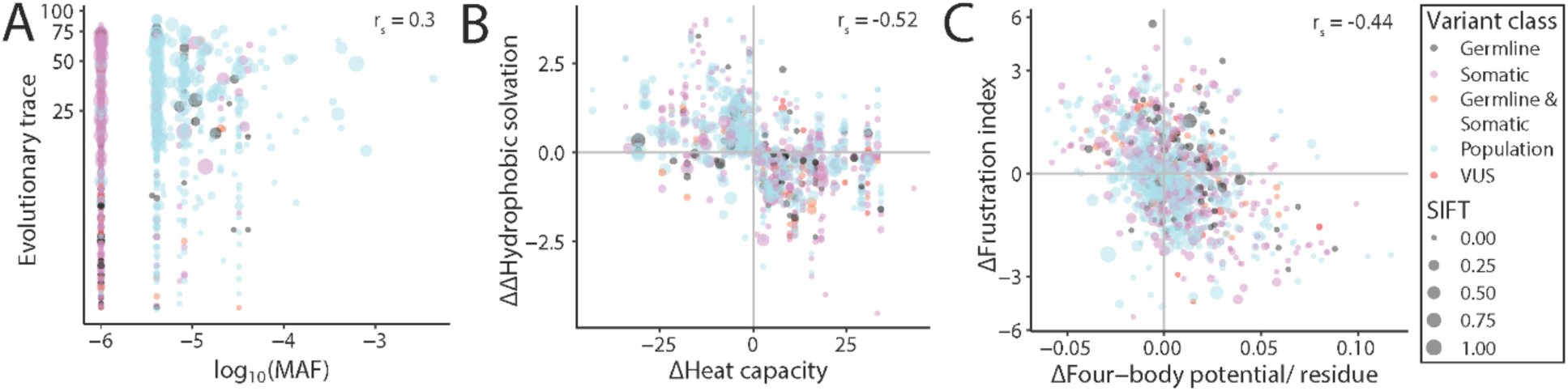
Patterns among scores. **A-C)** Patterns between pair of selected scores are shown as scattered plots for five variant classes indicated by different colors. The impact of amino-acid substitution on protein function are represented by the SIFT score as different sizes for the points in the scattered plots. Lower SIFT score (smaller point sizes) indicates higher predicted impact on protein function. Higher minor allele frequency (MAF) indicate the variants are more common in population. The larger the phyloP30way score, the more conserved the site. Negative change in the frustration index indicate increase in local energetic frustration (energetically less stable) upon mutation. Spearman correlation (r_s_) between the pair of scores are included inside each plot.

### An Approach for Integrating Molecular Scores that Predict Variant Classes and Mechanistic Clusters

Since individual scores or patterns among pair of scores do not clearly distinguish from among variant classes, we next combined all 31 scores (**Figure 1C and S4B**) and performed an integrated analysis using an unsupervised nonlinear dimensionality reduction process (t-SNE; see **Methods**) represented into a 2D space (**Figures 5A**). The t-SNE-generated dimensions (t-SNE1 and t-SNE2 in **Figures 5A**) are optimized in such a way that the scores of the variants, which are similar to one another in the raw high dimensional data (31 scores by 926 variants), are close in the 2D reduced data space. Importantly, t-SNE embeddings are non-Euclidian, making them difficult to quantify similarities among all samples, but helpful for inferring relative similarities. To better interpret the potential meaning of t-SNE dimensions, we superimposed the MAF in the t-SNE space (**Figure 5A**) to see the pattern in variant classes. We divided the t-SNE space in three topological groups (‘Group 1’, ‘Group 2’, and ‘Group 3’ in **Figure 5B**) under the hypothesis that variants within a topologic group will be more similar to each other in terms of likelihood of pathogenicity and underlying disease mechanism(s). We found that Group 1 is especially dominated by the variants that are likely to define diseases (Germline and Germline & Somatic variant classes), whereas Group 2 and Group 3 are dominated by the variants from Population and Somatic disease classes (**Table S3**). Thus, it appears that each topologic group in the t-SNE space likely reflects different levels of pathogenicity probability for RAS variants.

**Figure 5:**
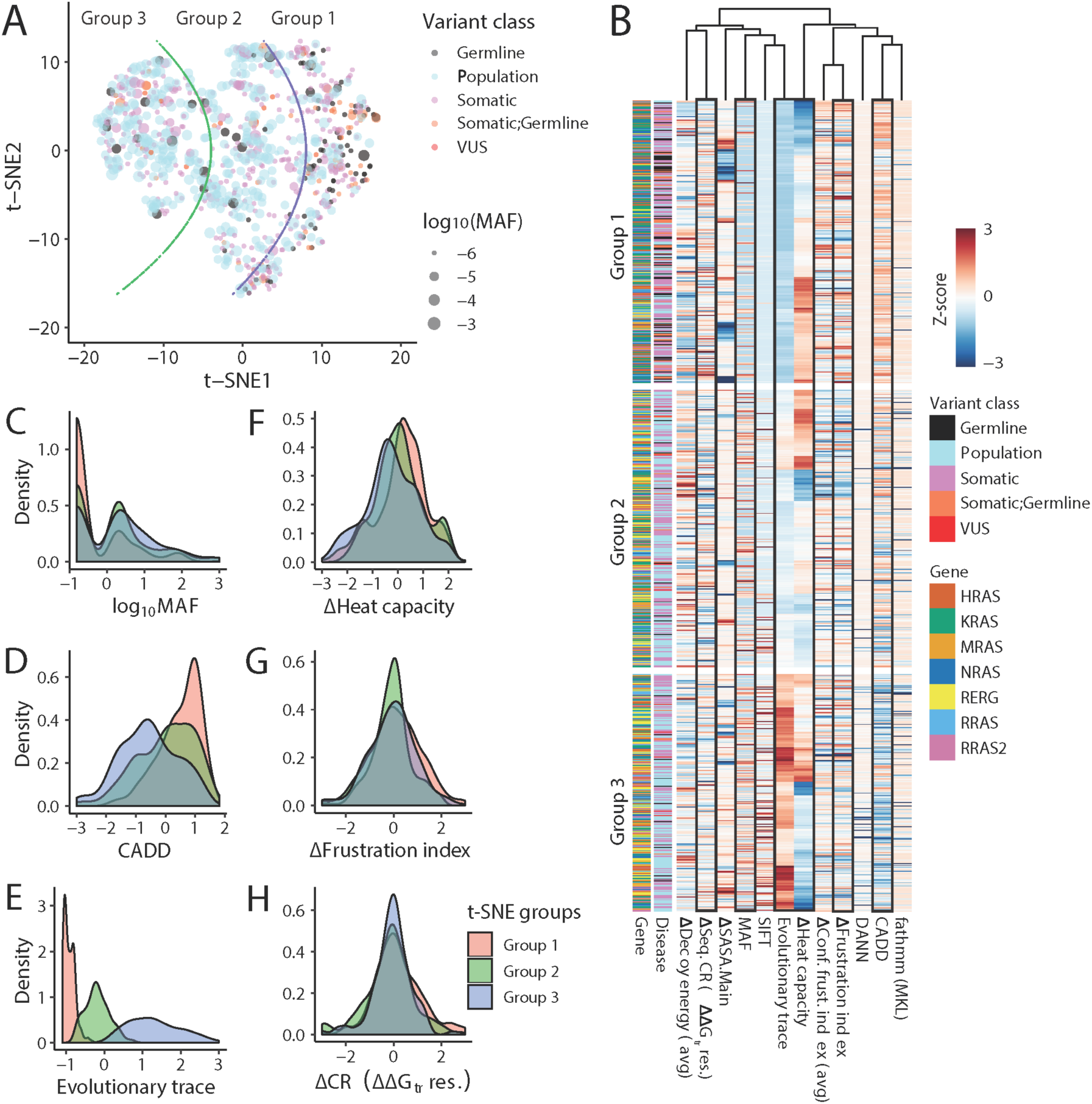
Combinations of scores better convey the similarities among variants and may indicate disease mechanisms. t-Distributed Stochastic Neighbor Embedding (t-SNE) was performed to reduce dimensionality into a 2D data space. **A)** Distribution of variants in the t-SNE space for five variant classes indicated by different colors. The different sizes of each point representing the variants denote minor allele frequency (MAF). Lower log_10_(MAF) values (smaller point sizes) indicate the variants are rare. Variants are divided in three groups (‘Group 1’, ‘Group 2’ and ‘Group 3’) based on the topology of the t-SNE space indicated by green and purple curves. **B)** Heatmap plot shows patterns of the top twelve scores selected based on one-way ANOVA cut-off (< 1×10^−4^) with t-SNE groups as factor. Heatmap plot with all scores are shown in **Figure S6**. **C-H)** Distributions of selected scores (indicated by black boxes in **B**) as z-scores for the t-SNE topologic groups.

We also explored the patterns of the scores in the t-SNE topological groups to see how each of them varies among these groups (**Figure S6**). As shown in **Figure 5B**, using ANOVA we identified 12 of the 31 scores that have the most distinct patterns among the t-SNE groups. Next, we focused on the 1D distribution of 6 scores from the abovementioned 12 scores that have the largest shift among t-SNE topological groups (**Figure 5C-H**). For MAF we observed population-shift among the t-SNE groups (**Figure 5C**). We observed a mean-shift in CADD (**Figure 5D**), ET (**Figure 5E**), and heat capacity (**Figure 5F**) scores among the t-SNE topological groups, whereas a variance-shift is observed in the difference in frustration index (**Figure 5G**) and the sequence-based CR difference (**Figure 5H**). Notably, using MAF, Group 1 is identified as having a greater number of very rare variants compared to Groups 2 and 3. We further observe greater CADD scores in t-SNE topologic Group 1 compared to Group 2 and Group 3 (**Figure S7**), indicating that Group 1 variants are more likely to alter protein function. More importantly, the ET score clearly separates the three t-SNE groups suggesting variants in Group 1 is highly conserved, Group 2 is moderately conserved, and Group 3 is not conserved (**Figure S7**). Thus, RAS variants in each topologic group (in the t-SNE space) most likely implicate distinct diseases mechanisms.

We analyzed missense gain-of-function variants in KRAS, HRAS and NRAS, which define the 3 distinct hotspot residues: G12, G13 and Q61 [6]. Notably, we find all variants affecting the three hotspot residues are not only part of Group 1 in the 2D t-SNE space but are also close to one another in the t-SNE space irrespective of the RAS gene (**Figure 6A-C**). We further notice that Q61 variants (**Figure 6C**) are more widely distributed in the t-SNE space compared to the G12 (**Figure 6A**) and G13 (**Figure 6B**) variants. From this pattern of the hotspots variants we hypothesize that certain hotspot variants have an effect that is the same across all RAS proteins, while others have a mechanism that is more distinct for each RAS protein.

**Figure 6:**
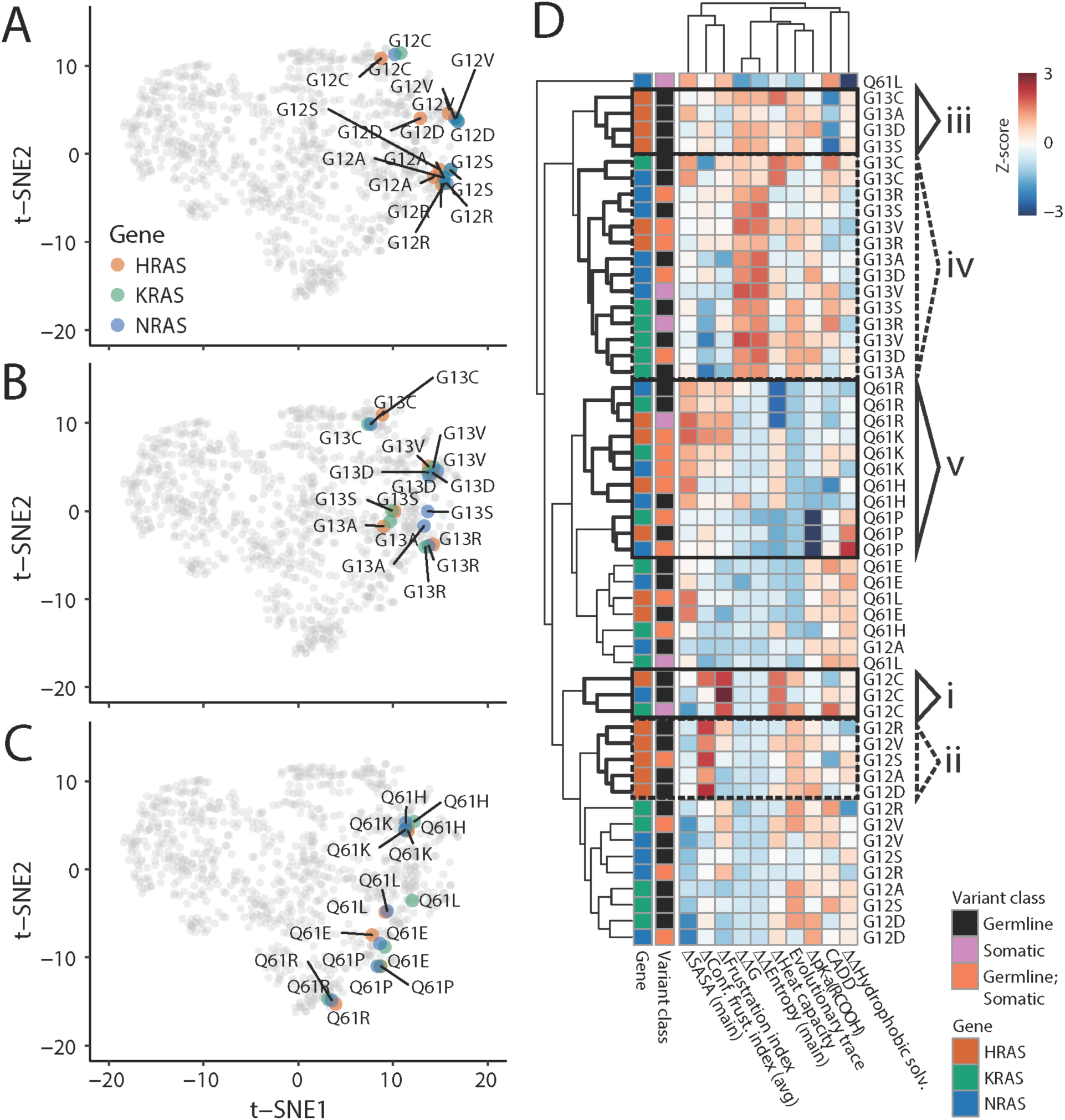
Mechanistic differences in the effect of different somatic hotspot variants. **A-C)** Somatic hotspot variants of G12 (**A**), G13 (**B**) and Q61 (**C**) are shown in the 2D t-SNE space for HRAS, KRAS and NRAS. **D)** Heatmap plot shows patterns of scores across t-SNE topologic groups for the somatic hotspot variants. The top ten scores are selected based on median absolute deviation criteria (≥ 0.2). Heatmap plot with all scores are shown in **Figure S8**. Specific regions in the heatmap are enclosed inside the solid (indicated as ‘i’, ‘iii’ and ‘v’) and dotted (indicated as ‘ii’ and ‘iv’) rectangles (see Main text).

Thereupon, we compared scores using a heatmap to see how their pattern varies among the same hotspot variants from different RAS and among different hotspot variants from the same RAS (**Figure 6D and S8**). First, we note that irrespective of the RAS gene all G12C variants are clustered together (indicated as group ‘i’ in **Figure 6D**). Other G12 variants (G12R, G12V, G12S, G12A and G12D) have a different pattern. Those from HRAS are clustered together (‘ii’ in **Figure 6D**), whereas those from KRAS and NRAS are clustered together. Similarly, the G13 variants G13C, G13A, G13D and G13S, belong to the same cluster (‘iii’ in **Figure 6D**) for HRAS, indicating that the HRAS context for these variants has a different effect than when they occur in KRAS or NRAS, while other G13 variants from HRAS, KRAS and NRAS are clustered together (‘iv’ in **Figure 6D**), indicating that the alternate amino acid has about the same effect in all three proteins. For Q61, the variants Q61R, Q61K, and Q61H are all clustered together across proteins (‘v’ in **Figure 6D**). This distinct pattern of the scores for the 3 hotspot RAS variants not only elucidates the sensitivity of each score but also the potential gains by integrating them for interpreting the impact of variants on RAS function.

### Evaluation of Integrated Molecular Scores for Variant Classification

We used the variant classes Germline (10.9% of the dataset), Population (43%), Germline & Somatic (5.7%) and Somatic (40.4%) for the prediction of function scores for each RAS variant. Thus, classification of the variants from the RAS family proteins is a multinomial classification problem and we note that there is an imbalance in the number of variants among the four classes as well as biologic overlap in their definition. Many Machine learning (ML) models pursue the goal of distinguishing disease-related variants from non-disease variants. However, it is unknown which somatic variants change function and which are not, or if variants observed in both germline and somatic diseases differ from those only observed in cancer. Therefore, we seek to provide guidance on the combination of scores that may be most informative for distinguishing among the effects of variants and if there are differences in the patterns of scores among these four classes.

ML identified MAF as the most important feature across all top models in the stacked ensemble (**Figure 7A-C and S9**). Protein 3D structure-based scores are also found to be important in the top ML models. Contribution from the change in mainchain entropy, main chain SASA, 4-body potential, ΔΔG_fold_, and sidechain-sidechain and sidechain-backbone hydrogen bonds are frequently the 3^rd^ through 5^th^ most important features across the ML models. Protein sequence-based conservation score is the second most important feature. DNA sequence-based conservation score fathmmMKL also shows importance across different ML models but is at most the 6^th^ highest-weighted feature. Thus, for predicting variant class, a combination of features from all three molecular levels provides the strongest performance.

**Figure 7:**
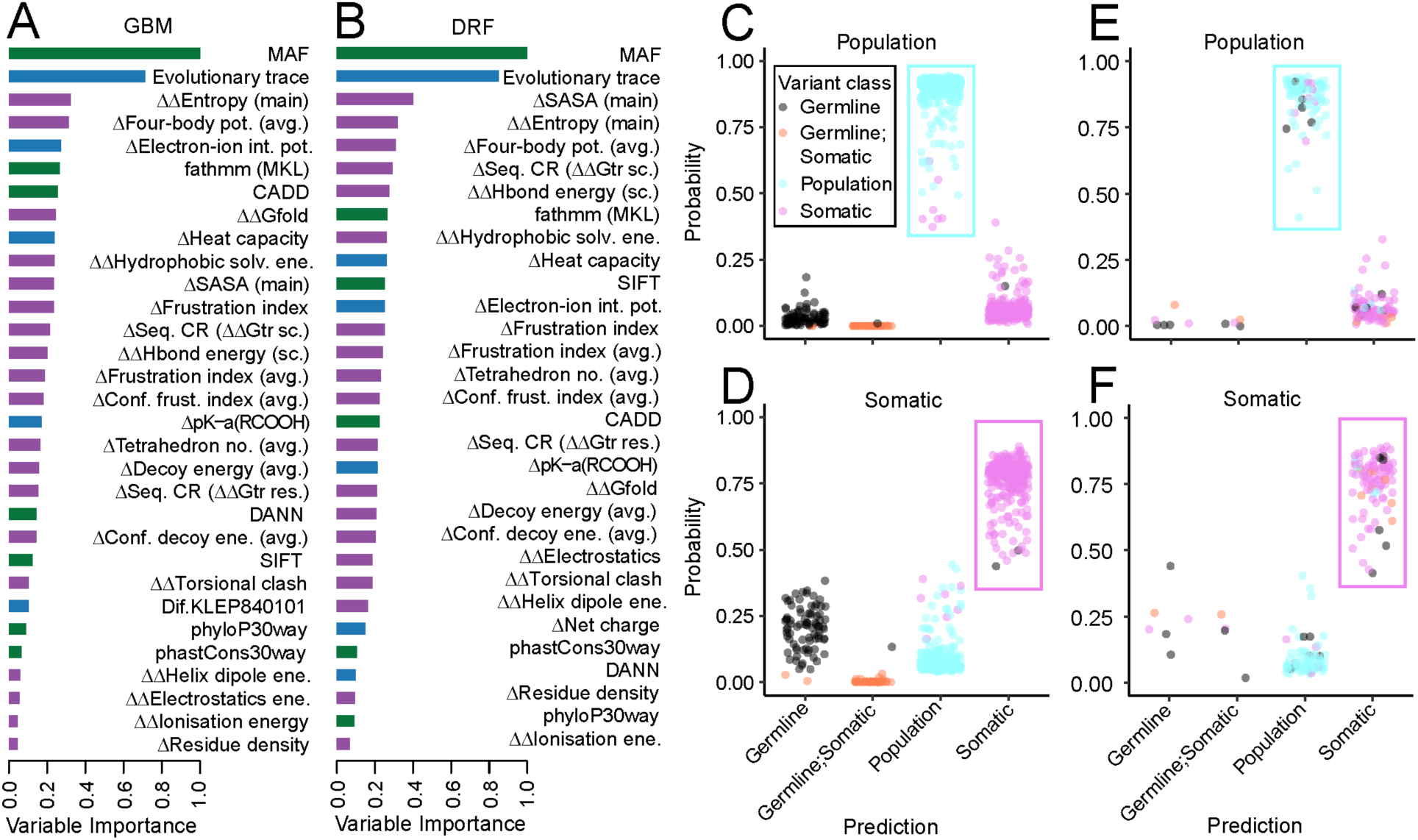
Machine learning optimally combines individual scores. **A-B)** Feature importance from two ML models from the stacked ensemble, gradient boosting machine (GBM) and distributed random forest (DRF), colored by different molecular levels as in **Figure 1**, demonstrate the importance of using scores from across all molecular levels. We abbreviated side chain as “sc.”, average as “avg.” See **Methods** for full details. Feature importance from all the ML models are shown in **Figure S9**. **C-F)** Associations for ML-emitted predictions of variant classification for the train set (**C, D**) and for the test set (**E, F**) for Population (cyan color rectangle in **C** and **E**) and Somatic (magenta color rectangle in **D** and **F**) variant classes.

Performance of the prediction of the variant classes on the training dataset showed high accuracy with only 1.63% overall error (**Table S4**). In fact, all the variants in the Population class are predicted correctly, whereas for the Germline & Somatic and Germline classes the error is 4.65% and 3.57%, respectively. Because there is likely biologic overlap among these classes, errors among these classes may be an indication of which variants share common effects even though they are observed in different disease contexts. On the other hand, overall accuracy for the prediction of the variant classes on the test data set is 19.89% (**Table S5**). Although, the Population and Somatic classes has error of 6.49% and 11.49%, respectively, the Germline & Somatic and Somatic classes show high error of 90% and 82.35%, respectively. We believe that imbalance in the variant numbers in each class and the systematically low MAF of the Germline & Somatic and Somatic classes are the reason for this high classification error, and also an indication that the effect on RAS for these variants may be, on average, similar. Therefore, we have used ML to identify that a combination of features from different molecular levels can be used to classify disease versus non-disease variants and to generate additional hypotheses about underlying molecular mechanism.

## Discussion

We have developed an efficient process for applying our multi-layered scoring approach and used it to evaluate all observed missense variants across 7 RAS-family proteins. We have identified patterns among these scores that can be used to group variants into their potential molecular mechanisms. Interestingly, our approach predicts that groups of individual variants at somatic hotspots may have different underlying molecular mechanisms, congruent with a small but growing amount of experimental evidence but for which no explanations are forthcoming. Thus, the need for more specific methods to interpret genomic variants is high, and our approach of adding 3D data has high potential to bring the specificity required for improving variant interpretation.

We have assembled a large and diverse group of scores from DNA annotations, protein sequence properties, and protein 3D properties. We showed that use of 3D scores enhances the information available from the DNA, leading to greater specificity. As a first example, several studies have shown that pair-wise energetic potentials can reliably identify changes in protein stability (ΔΔG_fold_) associated with genomic variants [15-17]. Second, due to allosteric transitions and biomolecular recognition sites, variants throughout the protein can lead to local functional changes without strongly disrupting global stability. In this context, local frustration quantifies the balance (or imbalance) among energetically favorable and unfavorable interactions [18] and has been shown useful to interpreting the impact of genomic variants [19]. Thirdly, four-body contact potentials were developed to identify the best 3D models from a pool of candidate models because they better capture the non-linear protein fold as well as the many interactions between the protein backbone, side chains, and solvent, compared to pair-wise potentials [20]. We further quantified an experimentally parameterized thermodynamic measure of local foldability using transfer free energies based on residue-specific implementation of the structure-energy-equivalence-of-domains (SEED) algorithm [21], which parses proteins into their constituent thermodynamically cooperative components [22]. Finally, we integrated scores and generated topologic groups that we believe may indicate different molecular mechanisms and thereby probabilities of pathogenicity, but more quantitative groups generated by training against experimental functional assays is required for a more definitive assessment. Earlier studies have shown the importance of protein 3D structure [5, 23, 24] and dynamics [4] in the prediction functional impact of missense variants, in general. However, no study, to our knowledge, has combined these diverse scores together, or with DNA annotations, for the interpretation of genomic variants. Thus, our approach is novel, generalizable to other proteins, and has a high potential to elucidate altered mechanisms.

As a first step towards a standardized and parametric approach to multi-level molecular scoring, in this study, we have investigated which properties predict the context each RAS variant was observed in from among germline disease, somatically in cancer, both, and neither. While there are four groups, they are not independent, and our main outcome is to distinguish disease from non-disease variation. However, which somatic variants are functional and have a greater or lesser effect than others are unknown. Thus, the ability to both categorize and grade the effect of genomic variants in greater specificity is needed. In particular, assessing the complexities of functional dysregulation is a critical step to surmounting the challenge of variant interpretation. The most well-studied RAS mechanism of dysregulation is hotspot variation, which are activating and commonly observed in cancer. A simple functional hypothesis is that all hotspot variants are damaging to function and activating, but an increasing array of evidence is indicating that different changes at the same hotspot residue result in different amounts of dysregulation or even different types of activation [25]. For example, in our data, G12C is more alike across RAS proteins than other G12 variants; our approach predicts that G12C affects RAS differently than other G12 variants. Recent experimental studies have shown that G12C is unique among G12 variants for its ability to be specifically inhibited [26]. Functional genomics experiments will be critical for completing our understanding of how mechanistic changes to RAS lead to distinct cellular effects. Concordance between existing experiments and predictions from 3D scores highlights the potential utility of our approach for identifying underlying mechanisms.

The primary aims of the current study were to quantify the difference in information content among DNA and 3D structure-based scores, and to investigate if there were groups of RAS variants based on how they alter the landscape of scores. We aim to define new scores that more directly assess biologic mechanisms of dysfunction. That is, to define energetic, parametric, or molecular mechanic scores that convey an underlying biophysical landscape or mechanism of alteration, even if they do not directly measure that landscape. Our long-term aim is to categorize variants into mechanistic groups. Such mechanisms are the underpinnings for disease. Thus, we aim to predict pathogenicity by first determining mechanism.

We have established our approach to protein structure-based scores as an initiating point for a fuller description for how genetic variants may affect protein function. Pilot studies on RAS proteins [14, 27-29], and our own studies on other proteins, [30-34] have demonstrated the utility of atomic molecular simulations to provide additional information such as allosteric transitions and functional motion, but scalable approaches using these tools remain to be developed. Our ongoing work will extend the structure-based approach presented here, to include dynamics-based scores, as well as scores derived from protein shape and surface properties. We will extend our application of protein scores to include the GTPase fold in general and further details for variants determining RASopathies. We firmly believe that the approach we have presented here is applicable to a broad range of the human proteome and will become an important criterion in future versions of guidelines for the interpretation of genomic variants.

## Conclusions

DNA sequence-based information is the mainstay of genomics data interpretation, but we have demonstrated in this work that additional information relevant for interpreting genomic variants and not currently predictable or available from the DNA sequence can be derived from computational study of the protein 3D structure. Within 3D analyses, there are multiple types of features that indicate alterations in different properties of RAS proteins, and likely proteins in general. Therefore, assaying one type of feature is insufficient for genomics data interpretation. A full suite of scores that integrate across all biologic layers of molecular function is required. This study demonstrates the potential for genomic data interpretation, that such an integrated system provides.

## Methods

### Gene Annotation

We used data from Orphanet [35] and Monarch [36] databases to identify mappings between rare diseases and RAS family genes.

### Genomic Variant Annotation

We first defined the GTPase domains of seven RAS family proteins in 3D and identified their corresponding DNA coding regions in the genome for UniProt canonical isoforms. We used COSMIC [37] to identify genetic variants previously observed somatically in human cancers. We used ClinVar [38] and HGMD [39] to identify genomic variants responsible for congenital diseases. We identified variants observed exclusively in the currently healthy adult population, and the global minor allele frequency (MAF) of disease variants from gnomAD [40].

Because of codon degeneracy, multiple DNA changes could result in the same amino acid change. In these cases, the protein change is the same, but scores from the DNA may differ from each other. Therefore, to be conservative, we chose the most severe score of the variants observed within the codon. Annotations for DNA sequence-based scores were gathered from dbNSFP v4.0 [41]. We used the BioR v5.0.0 [42] system for handling genomic data resources with custom scripts in the R programming language [43] for analysis.

We defined pathogenic variants as any genomic variant from ClinVar or HGMD annotated as (likely) pathogenic and lacking conflicting annotations among either database. We defined benign variants similarly. Functional variants from ClinVar, such as drug-response phenotypes, that are also considered pathogenic in HGMD were considered pathogenic variants. All other variants from ClinVar and HGMD were considered VUS. We defined polymorphic variants as those from gnomAD with MAF ≥ 1%. GnomAD is reflective of the populations sequenced, while protein structure is applicable to either germline or somatic setting and protein structural data are independent of the germline frequencies. Future sequencing work to be more representative of the global population and thereby closer to an objective property of the human genome, while knowledge gained about each protein variant will remain.

### Disease Categories of Genomic Variants

Using these resources, we classified the disease context of each genomic variant. Variants known to cause germline diseases are referred to as “Germline.” Variants observed somatically in cancer and lacking any association with a defined germline phenotype are referred to as “Somatic.” Variants that have both a (likely)pathogenic role in a germline setting and are also observed somatically in cancer and referred to as “Germline & Somatic.” Variants that do not have any association with germline or somatic disease, but are observed in the currently healthy adult population, are referred to as “Population.” Finally, all genomic variants that do not fit one of these categories, or for which the clinical annotations are conflicting, are considered variants of uncertain significance (VUS).

### Molecular Modeling

GDP-bound structures were used for all the RAS proteins in this study (**Table S1**). To model the missing loops in NRAS (PDB: 3CON, residue 61–71) and RRAS2 (PDB: 2ERY, residue 71–74 in chain A) ModWeb (https://modbase.compbio.ucsf.edu/modweb/) server was used.

### Protein Variant Scoring

For RAS missense variants, we combined scores at different molecular levels based on sequence (DNA and protein), amino-acid properties, and 3D protein structure using several available tools. These tools do not explicitly include ligand-protein interactions but focus on evaluation of the naturally occurring amino-acids. Correlation between all the scores are shown in **Figure S4A**.

#### DNA sequence-based scores

The sequence-based scores Sorting Intolerant From Tolerant (SIFT) [44], Polymorphism Phenotyping v2 (Polyphen2) [45], Meta-analytics Support Vector Machine (MetaSVM) and Logistic Regression (MetaLR) [46], Rare Exome Variant Ensemble Learner (REVEL) [47], Combined Annotation-Dependent Depletion (CADD) [48], Deleterious Annotation of genetic variants using Neural Networks (DANN) [49], Functional Analysis Through Hidden Markov Models based on multiple kernel learning (FATHMM-MKL) [50, 51] and with eXtended Features (FATHMM-XF) [52], phylogenetic P-value conservation score based on 30way alignment mammalian set (phyloP30way_mammalian and phastCons30way_mammalian) were extracted from dbNSFP v4.0 database [41].

#### Protein sequence-based scores

We calculated the degree of conservation of each amino-acid of RAS proteins using Universal Evolutionary Trace server (http://lichtargelab.org/software/uet). The real-value Evolutionary Trace (rvET) scores for each residue were calculated based on the phylogenetic tree among RAS sequences. This method assumes that the more functionally important the residue, the sooner it becomes fixed in the evolutionary tree [53].

#### Amino-acid physicochemical properties-based scores

For each missense variant we computed the change in physicochemical and biochemical properties upon amino-acid mutation based on the information in the AAindex database [54] derived from published literature for each variant using BioSeqClass R-package [55]. We evaluated all AAindex properties (**Figure S10**) and selected a subset for further analysis based on Spearman correlation and biochemistry, including electron interaction potential (COSI940101), polarizability (CHAM820101), pK-a(RCOOH) (FAUJ880113), net charge (KLEP840101), heat capacity (HUTJ700101), melting point (FASG760102), hydrophobicity (JURD980101), and normalized flexibility parameters (VINM940101).

#### Structure-based scores

We estimated the change in protein stability upon amino-acid substitution by calculating the change in folding free-energy, ΔΔG_fold_ (total.energy) using FoldX [56] for each variant; FoldX terminology is included in parentheses. Other energy parameters which we included from FoldX are the contributions of backbone hydrogen bonds (Backbone.Hbond), sidechain-sidechain and sidechain-backbone bonds (Sidechain.Hbond), VanderWaals (Van.der.Waals), electrostatic interactions (Electrostatics), penalization for burying polar groups (Solvation.Polar), hydrophobic groups (Solvation.Hydrophobic), penalization due to VanderWaals’ clashes (interresidue) (Van.der.Waals.clashes), entropy cost of fixing the side chain (entropy.sidechain), entropy cost of fixing the main chain (entropy.mainchain), VanderWaals’ torsional clashes (intraresidue torsional.clash), backbone-backbone VanderWaals (backbone.clash; not included in ΔΔG_fold_), electrostatic contribution of the helix dipole (helix.dipole), and contribution of the ionisation energy (energy.Ionisation). We used the FoldX generated structure of each variant for the evaluation of other structure-based scores.

To identify whether the solvent-accessible surface area (SASA) of side chains changed upon amino-acid substitution, we calculated absolute SASA for each variant of the RAS proteins using the NACCESS program [57] for all atoms (SASA.All), non-polar sidechain (all non-oxygens and non-nitrogens in the sidechain) (SASA.Nonpolar), polar sidechain (all oxygens and nitrogens in the sidechain) (SASA.Polar), total sidechain (SASA.Side), and mainchain (SAA.Main).

We computed ‘local energetic frustration’ in the RAS proteins for each variant using the Ferreiro-Wolynes algorithm [18, 58, 59], which estimates how favorable a specific contact (or an amino-acid residue) is relative to the set of all possible contacts (or an amino-acid residue) in that location and compares it to the energies of a set of ‘decoy’ states. All 20 amino acids are considered at each site in RAS for the calculation of frustration. For single residue level frustration (SRLF), decoys are constructed from the mutations of single residue by randomly selecting the amino acid identities in the native state. The local SRLF index (FrstIndex) for each amino acid is defined as a Z-score of the native state energy (NativeEnergy) compared to the N decoys (DecoyEnergy). Besides SRLF we also calculated mutational and configurational frustration at the residue-residue contact level. Mutational frustration (MF) measures how favorable are the native interactions between two residues relative to other residues in that location. On the other hand, configurational frustration (CF) measures how favorable are the native interactions between two residues relative to other interactions these residues can form in other compact structures. We converted the MF and CF to the residue level frustration by averaging over the number of contacts (Contact.no) for each residue, Mut_FrstIndex_Avg and Conf_FrstIndex_Avg, respectively. Additional parameters we included associated with SRLF are standard deviation in energy (SDEnergy) and local residue density (DensityRes) at the residue level, and parameters such as, NativeEnergy_Avg, DecoyEnergy_Avg, SDEnergy_Avg and FrstIndex_Avg, by averaging over all the residues in a protein. Similarly, for the MF and CF we calculated additional paramters such as, Mut_NativeEnergy_Avg, Mut_DecoyEnergy_Avg, Mut_SDEnergy_Avg and Conf_NativeEnergy_Avg, Conf_DecoyEnergy_Avg, Conf_SDEnergy_Avg. A contact is defined as ‘minimally frustrated’ if its native energy is at the lower end of the distribution of decoy energies, having a frustration index as measured with a Z-score of 0.78 or higher magnitude [18], that is, the majority of other amino acid pairs in that position would be energetically unfavorable. In contrast, a contact is defined as ‘highly frustrated’ if decoy energy is at the higher end of the distribution with a local frustration index lower than -1, that is, most other amino acid pairs at that location would be more favorable energetically for folding than the native ones by more than one standard deviation of that distribution. If the native energy is in between these limits, the contact is defined as ‘neutral’.

To characterize the difference in regional folding cooperativity between the wild-type and each variant we employed a residue-specific implementation of structure-energy-equivalence-of-domains (SEED) algorithm [21]. SEED parses proteins of known structure into their constituent thermodynamically cooperative components using residue-specific water ? 1M urea-group transfer free energies [22] to define thermodynamic subdomains of protein structures consistent with experimentally determined equilibrium folding intermediates. For each variant we computed the cooperativity ratio using sequential parsing for the amino-acid backbone (Seq_CR_DDGtr.BB), residue (Seq_CR_DDGtr.R) and sidechain (Seq_CR_DDGtr.SC).

Multi-body statistical contact potentials consider solvation-dependent three-dimensional interactions related to residue packing and capture the cooperativity of these interactions in protein structures. In this study, we calculated four-body contact potential (per residue; Pot.4body_res) and number of tetrahedrons (per residue; Tetra_res) by combining the four-body sequential with the four-body non-sequential potentials and with a short-range potential [20] from the 3D-strucutre of each variant.

### Grouping Variants by Similarity Across Scores

Analysis using t-distributed stochastic neighbor embedding (t-SNE) construct a low-dimensional embedding of high-dimensional data, distances or similarities. We used the Rtsne package v0.15 [60] based on optimized threshold estimate for a trade-off between speed and accuracy in a Barnes-Hut implementation of t-SNE [61] and a perplexity factor of 80 and theta is set to 0.3.

We then used machine learning to identify trends in the scores that could predict the context in which the variant has been observed. Data were standardized (z-score) prior to classifier input. We used H2O.ai’s AutoML v3.24.0.5 implemented in R v3.5.1 [43] for multi-classification of the variant class type. We divided our data (926 variants by 31 features) as 80:20 for the train and test sets, respectively. We used 20 models in H2O AutoML and “misclassification” as the stopping metric. StackedEnsemble_BestOfFamily_AutoML was identified as the model with best overall performance.

## Acknowledgements

This research was completed in part with computational resources and technical support provided by the Research Computing Center at the Medical College of Wisconsin.

## Tables

**Table S1:** Experimental protein structures used in this study.

| Protein | PDB ID |
| --- | --- |
| KRAS | 4OBE |
| HRAS | 4Q21 |
| NRAS | 3CON |
| RRAS | 2FN4 |
| MRAS | 1X1R |
| RERG | 2ATV |
| RRAS2 | 2ERY |
| HRAS Active | 3K8Y |
| HRAS Inactive | 4EFL |
| HRAS:GEF | 1BKD |
| HRAS:PI3K | 1HE8 |
| HRAS:GAP | 1WQ1 |
| KRAS:DARPIN K27 | 5O2T |

**Table S2:**
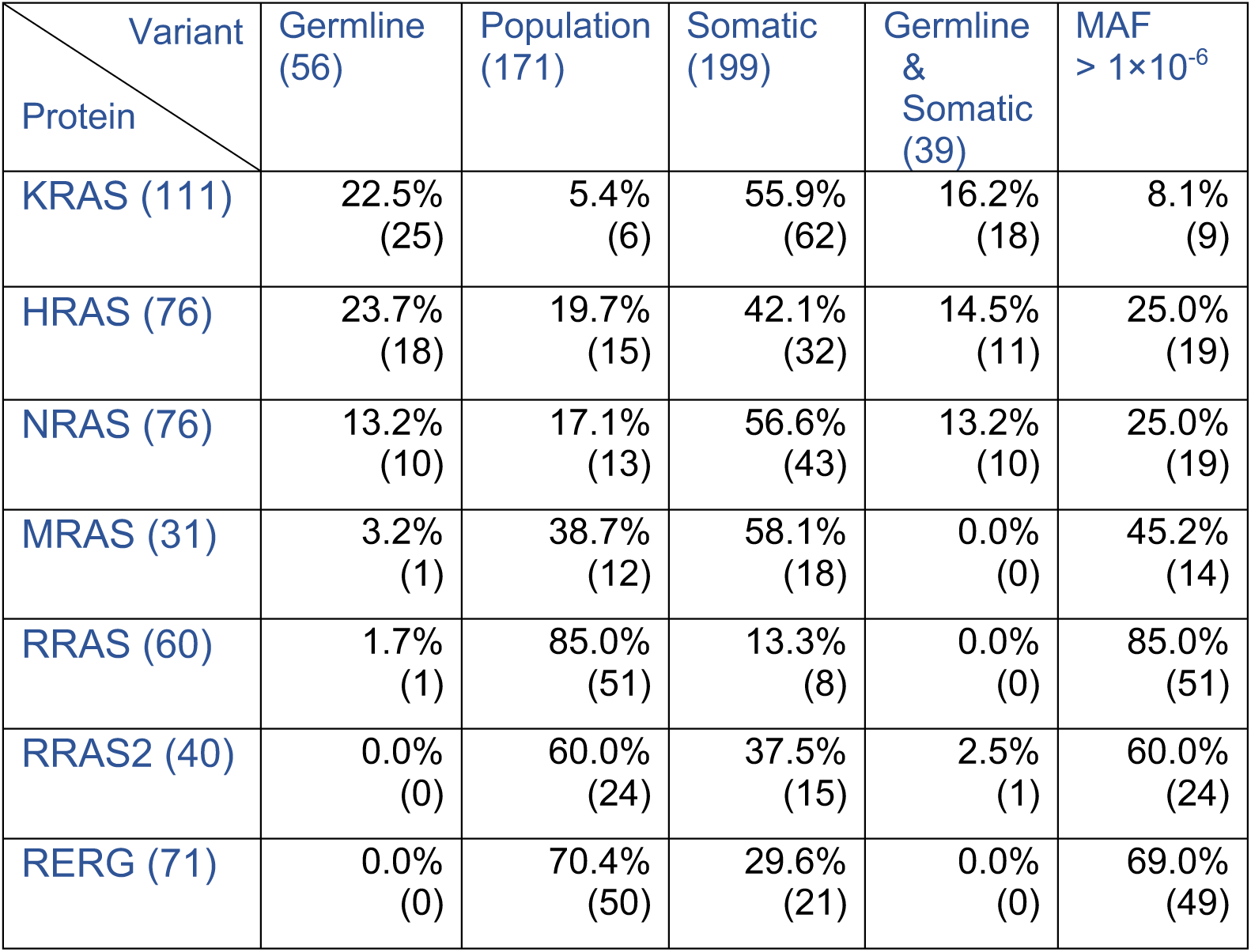
Distribution of 935 variants in the protein interface of 7 RAS proteins by variant classes (Supplemental data file).

| Variant<br>Protein | Germline<br>(56) | Population<br>(171) | Somatic<br>(199) | Germline<br>&<br>Somatic<br>(39) | MAF<br>> $1 \times 10^{-6}$ |
| --- | --- | --- | --- | --- | --- |
| KRAS (111) | 22.5%<br>(25) | 5.4%<br>(6) | 55.9%<br>(62) | 16.2%<br>(18) | 8.1%<br>(9) |
| HRAS (76) | 23.7%<br>(18) | 19.7%<br>(15) | 42.1%<br>(32) | 14.5%<br>(11) | 25.0%<br>(19) |
| NRAS (76) | 13.2%<br>(10) | 17.1%<br>(13) | 56.6%<br>(43) | 13.2%<br>(10) | 25.0%<br>(19) |
| MRAS (31) | 3.2%<br>(1) | 38.7%<br>(12) | 58.1%<br>(18) | 0.0%<br>(0) | 45.2%<br>(14) |
| RRAS (60) | 1.7%<br>(1) | 85.0%<br>(51) | 13.3%<br>(8) | 0.0%<br>(0) | 85.0%<br>(51) |
| RRAS2 (40) | 0.0%<br>(0) | 60.0%<br>(24) | 37.5%<br>(15) | 2.5%<br>(1) | 60.0%<br>(24) |
| RERG (71) | 0.0%<br>(0) | 70.4%<br>(50) | 29.6%<br>(21) | 0.0%<br>(0) | 69.0%<br>(49) |

**Table S3:** Distribution of variants in the t-SNE topological groups and variant classes.

| Variant<br>class<br>t-SNE<br>groups | Germline<br>(83) | Population<br>(400) | Somatic<br>(374) | Germline &<br>Somatic<br>(53) | VUS<br>(16) |
| --- | --- | --- | --- | --- | --- |
| Group 1<br>(328) | 72.3%<br>(60) | 19.0%<br>(76) | 39.3%<br>(147) | 77.4%<br>(41) | 25.0%<br>(4) |
| Group 2<br>(322) | 13.2%<br>(11) | 42.5%<br>(170) | 34.2%<br>(128) | 15.1%<br>(8) | 31.2%<br>(5) |
| Group 3<br>(276) | 14.5%<br>(12) | 38.5%<br>(154) | 26.5%<br>(99) | 7.5%<br>(4) | 43.7%<br>(7) |

**Table S4:** Confusion matrix for the train set of the concordant classes.

| Predicted class<br>Actual class | Germline | Germline & Somatic | Population | Somatic | Error | Rate |
| --- | --- | --- | --- | --- | --- | --- |
| Germline | 81 | 1 | 0 | 2 | 0.0357 | = 3/84 |
| Germline & Somatic | 2 | 41 | 0 | 0 | 0.0465 | = 2/43 |
| Population | 0 | 0 | 321 | 0 | 0.0000 | = 0/321 |
| Somatic | 0 | 0 | 7 | 280 | 0.0244 | = 7/287 |
| Totals | 83 | 42 | 328 | 282 | 0.0163 | = 12/735 |

**Table S5:** Confusion matrix for the test set of the concordant classes.

| Predicted class<br>Actual class | Germline | Germline & Somatic | Population | Somatic | Error | Rate |
| --- | --- | --- | --- | --- | --- | --- |
| Germline | 3 | 2 | 6 | 6 | 0.8235 | = 14/17 |
| Germline & Somatic | 1 | 1 | 0 | 8 | 0.9000 | = 9/10 |
| Population | 0 | 0 | 72 | 5 | 0.0649 | = 5/77 |
| Somatic | 2 | 1 | 7 | 77 | 0.1149 | = 10/87 |
| Totals | 6 | 4 | 85 | 96 | 0.1989 | = 38/191 |

## Supplemental Information

**Figure S1:**
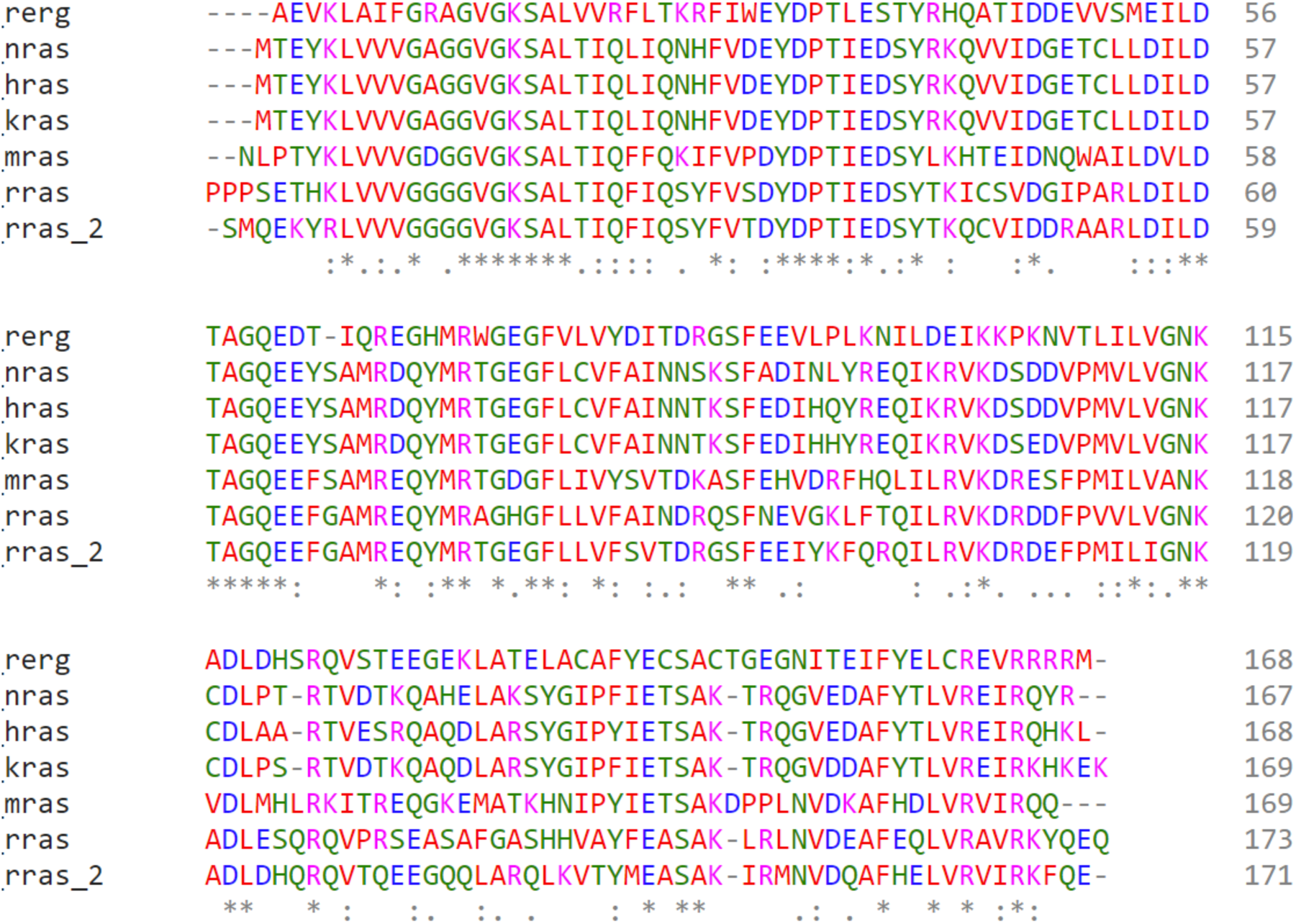
Multiple sequence alignment of RAS family proteins used in this study. Sequences are from the Protein Data Bank (PDB) structures (KRAS: 4OBE; HRAS: 4Q21; NRAS: 3CON; RRAS: 2FN4; MRAS: 1X1R; RERG: 2ATV; RRAS2: 2ERY) as indicated in **Table S1**.

**Figure S2:**
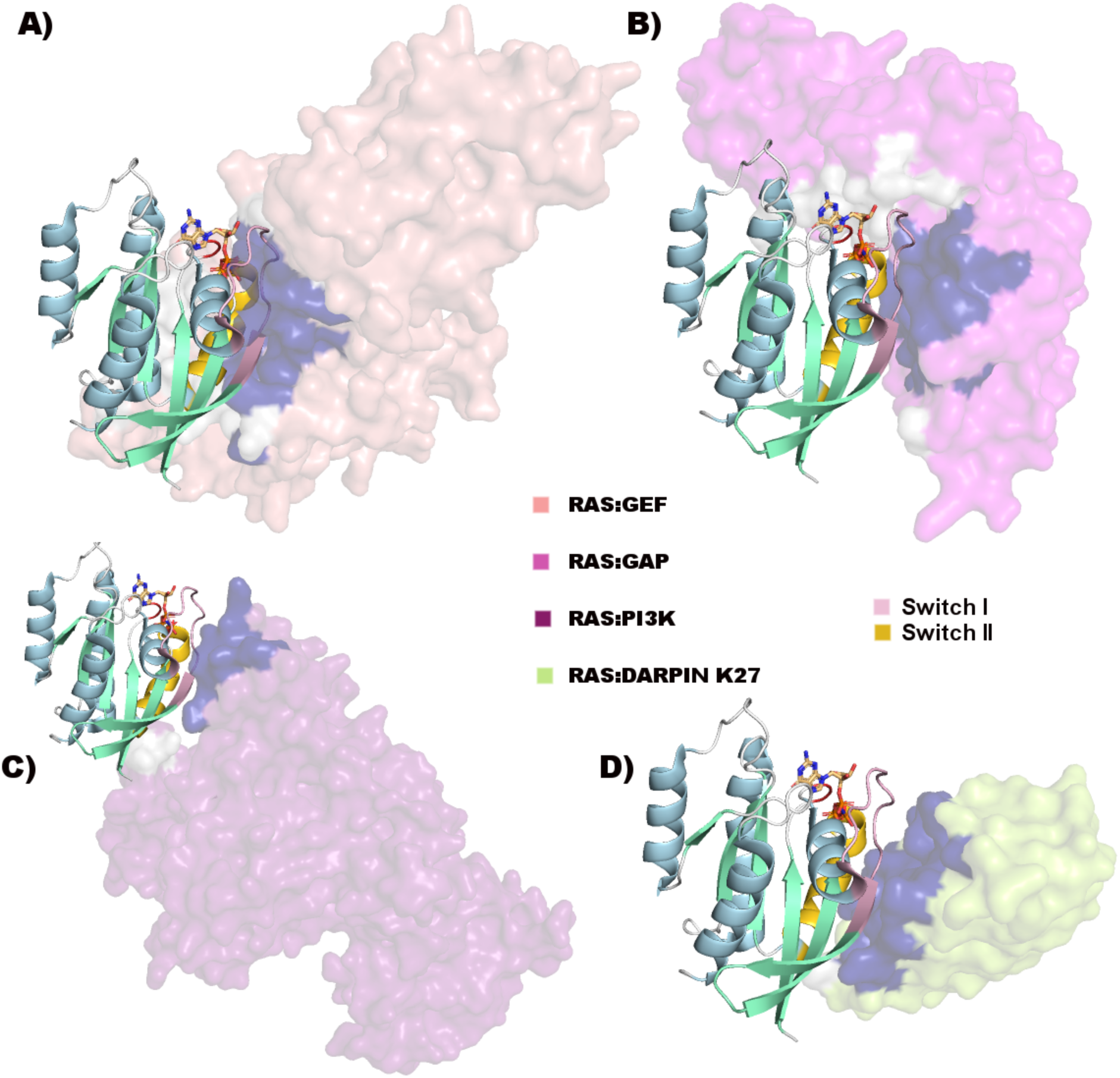
Structure of RAS proteins and their functional interactions. There are multiple experimental structures of each RAS protein, many of which include proteins whose interaction with RAS is necessary for specific functions (transparent surfaces). We show the interaction with **A)** GEF, **B)** GAP, **C)** PI3K, and **D)** darpin. Each interacting protein is colored distinctly, but with the surface contacting RAS colored blue or white for, respectively, interactions that are shared by all four interactions and those that are specific to a subset of the interactions. RAS is colored as in **Figure 2A**.

**Figure S3:**
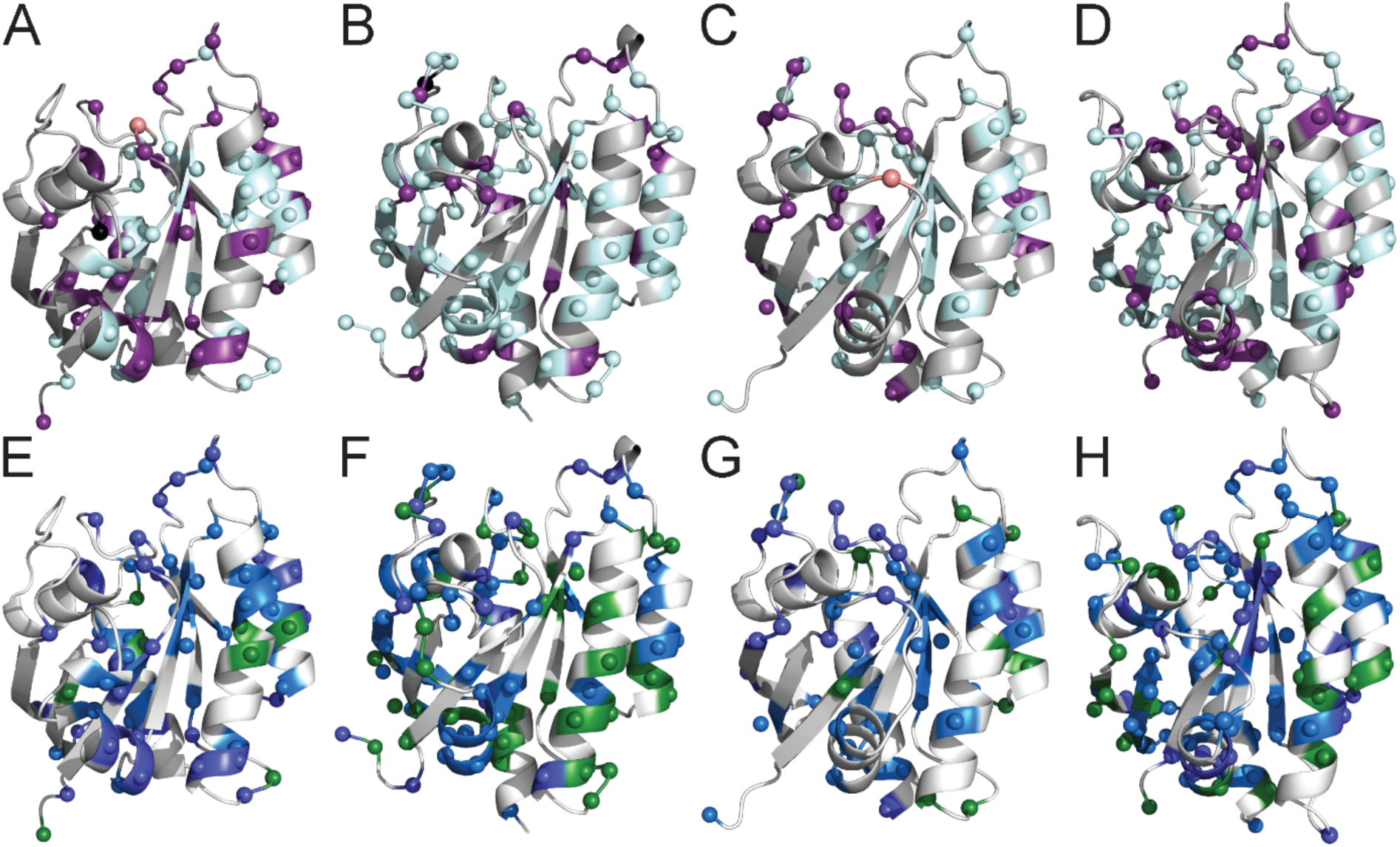
Structure of MRAS, RRAS, RRAS2 and RERG with variant classes (A-D, respectively) and MAF (E-H). Visualized as in **Figure 2**.

**Figure S4:**
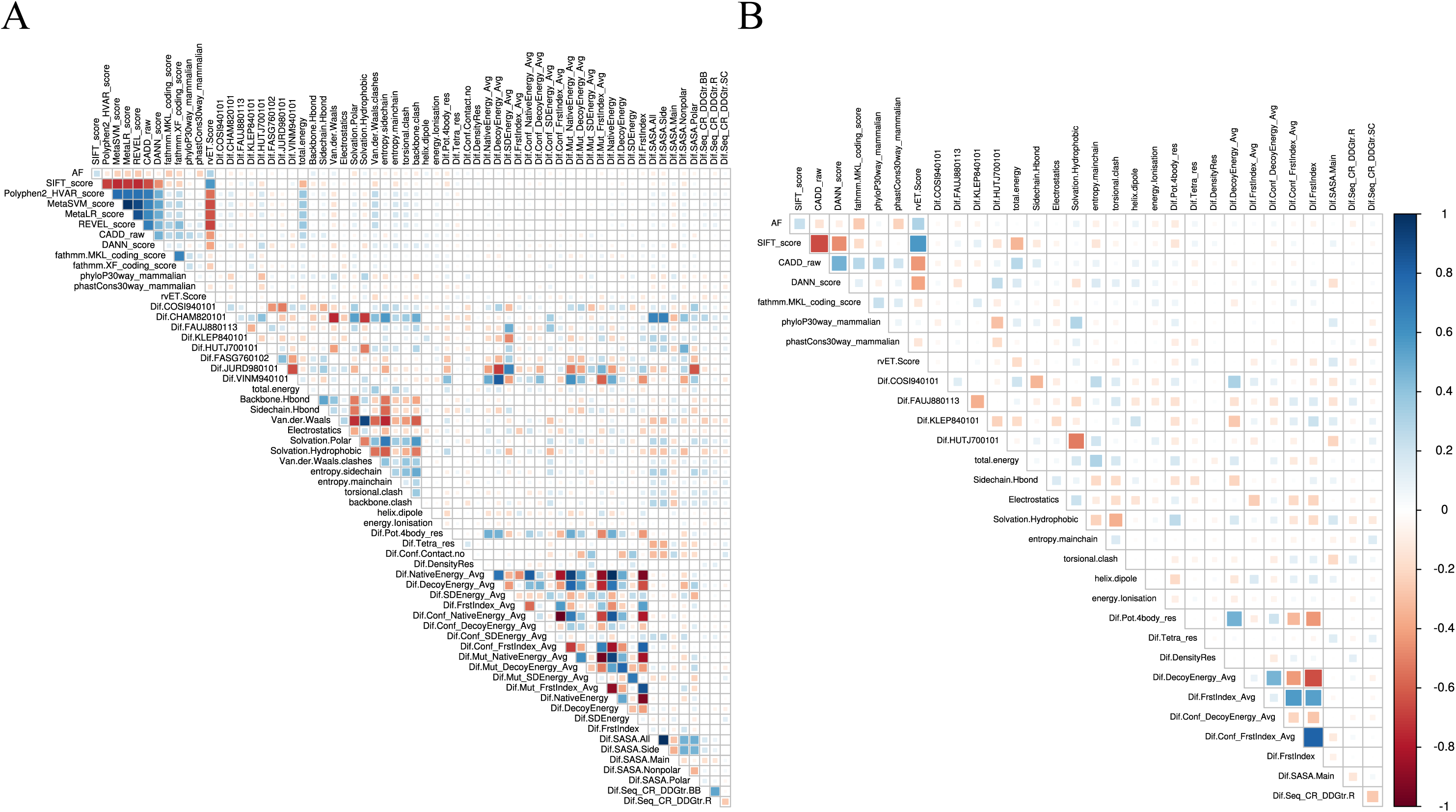
Choosing representative scores by domain knowledge and data correlation patterns. Spearman correlation data for the computed value of each protein variant subtracted from the corresponding WT value. **A)** The Spearman correlation matrix among 63 selected scores that capture information present in genomic DNA annotations, protein sequence properties, and protein structural features. These features were manually selected from many hundreds of candidate features using domain knowledge and literature precedence for best-in-class and unique measures of protein properties. We examined the correlation structure to further refine the set of granular scores by choosing a representative from pairs of scores that have a high absolute correlation. **B)** This procedure resulted in a final filtered selection of 31 granular scores.

**Figure S5:**
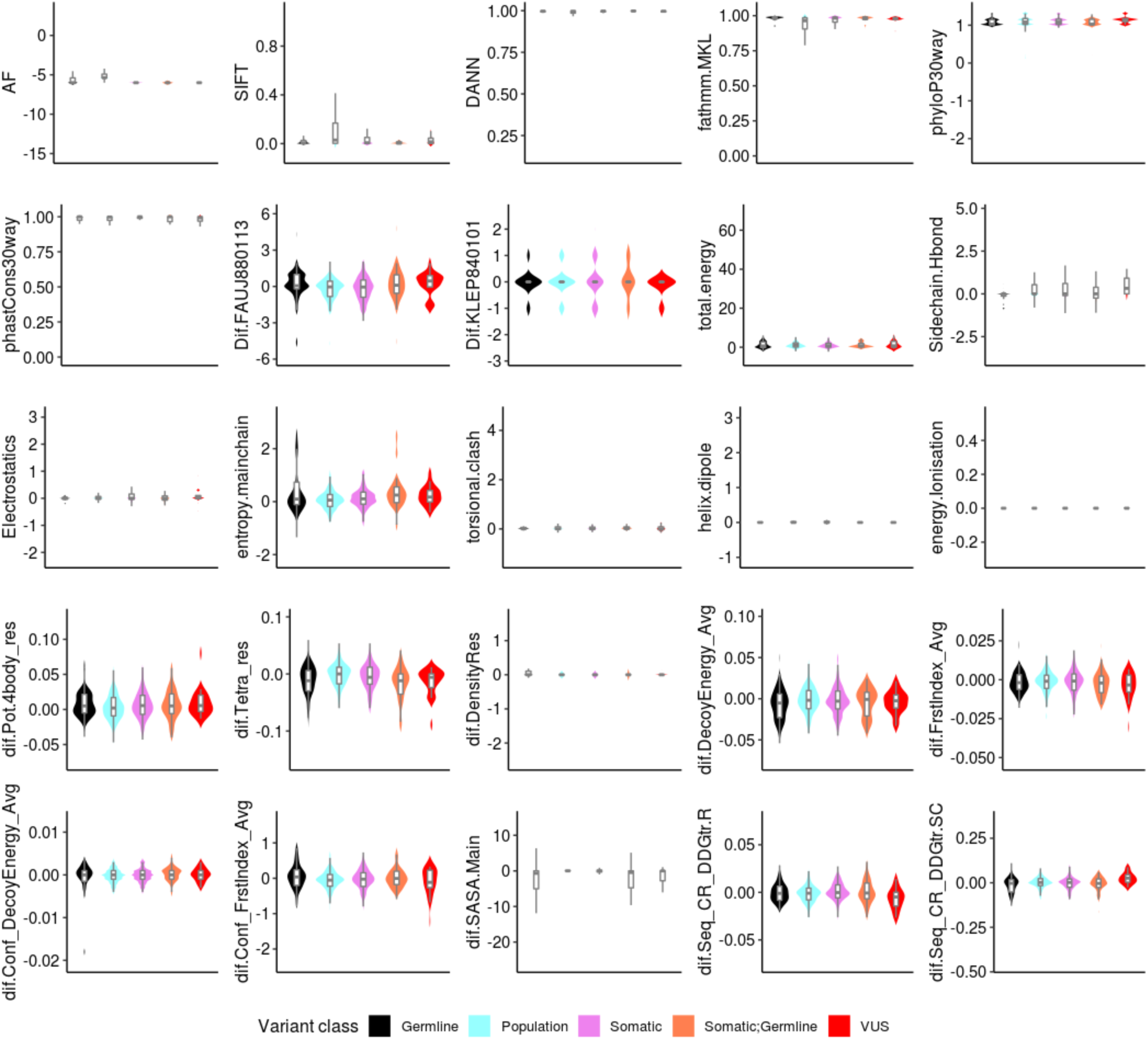
Distributions of 25 scores (out of 31) at different molecular levels are shown in violin plots for five variant classes indicated by different colors. The range of each score is set individually. Outliers are omitted from each plot. Distribution of six other scores are shown in **Figure 3**.

**Figure S6:**
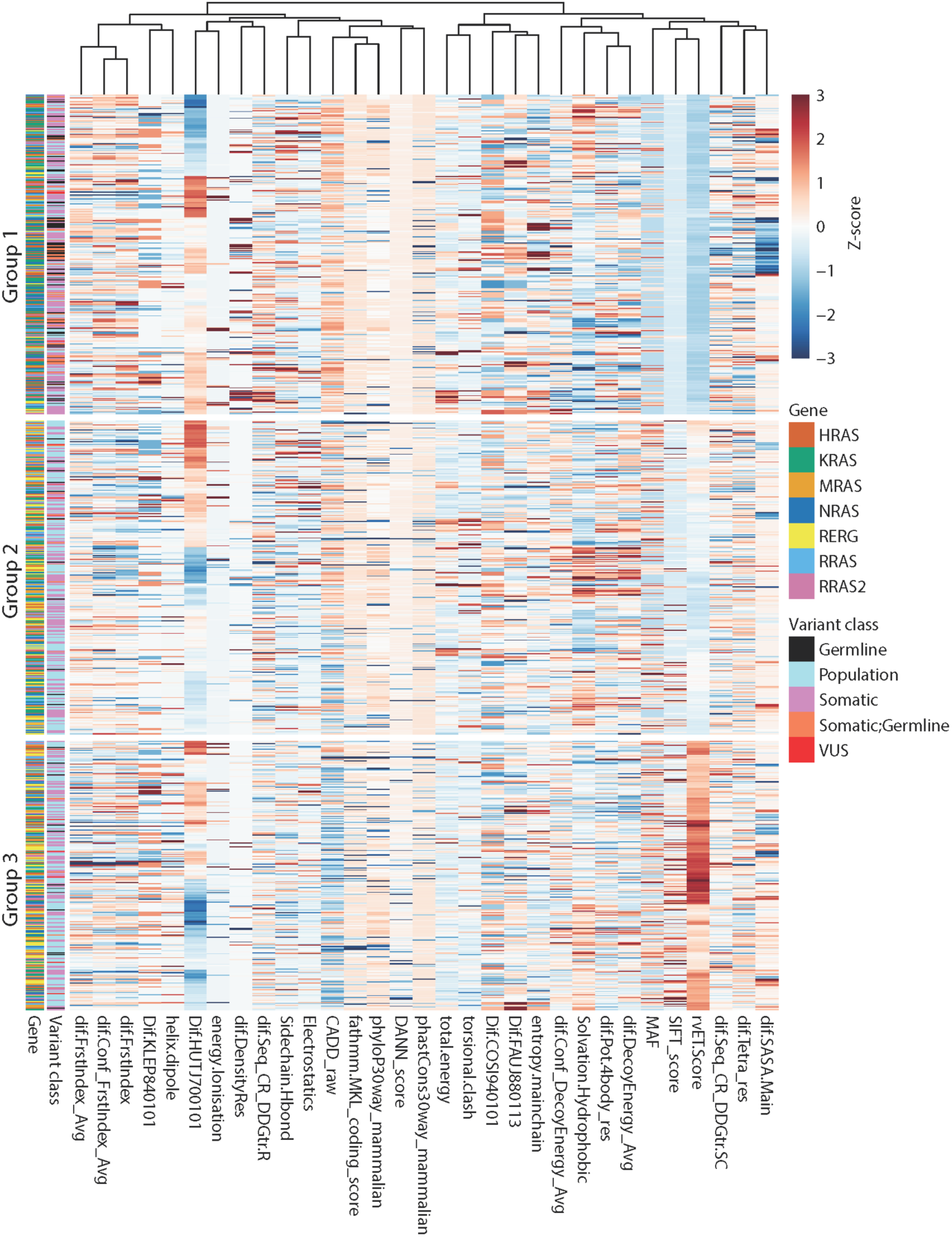
Heatmap plot shows patterns of scores across t-SNE topologic groups. Heatmap plot of selected scores are shown in **Figure 5B**.

**Figure S7:**
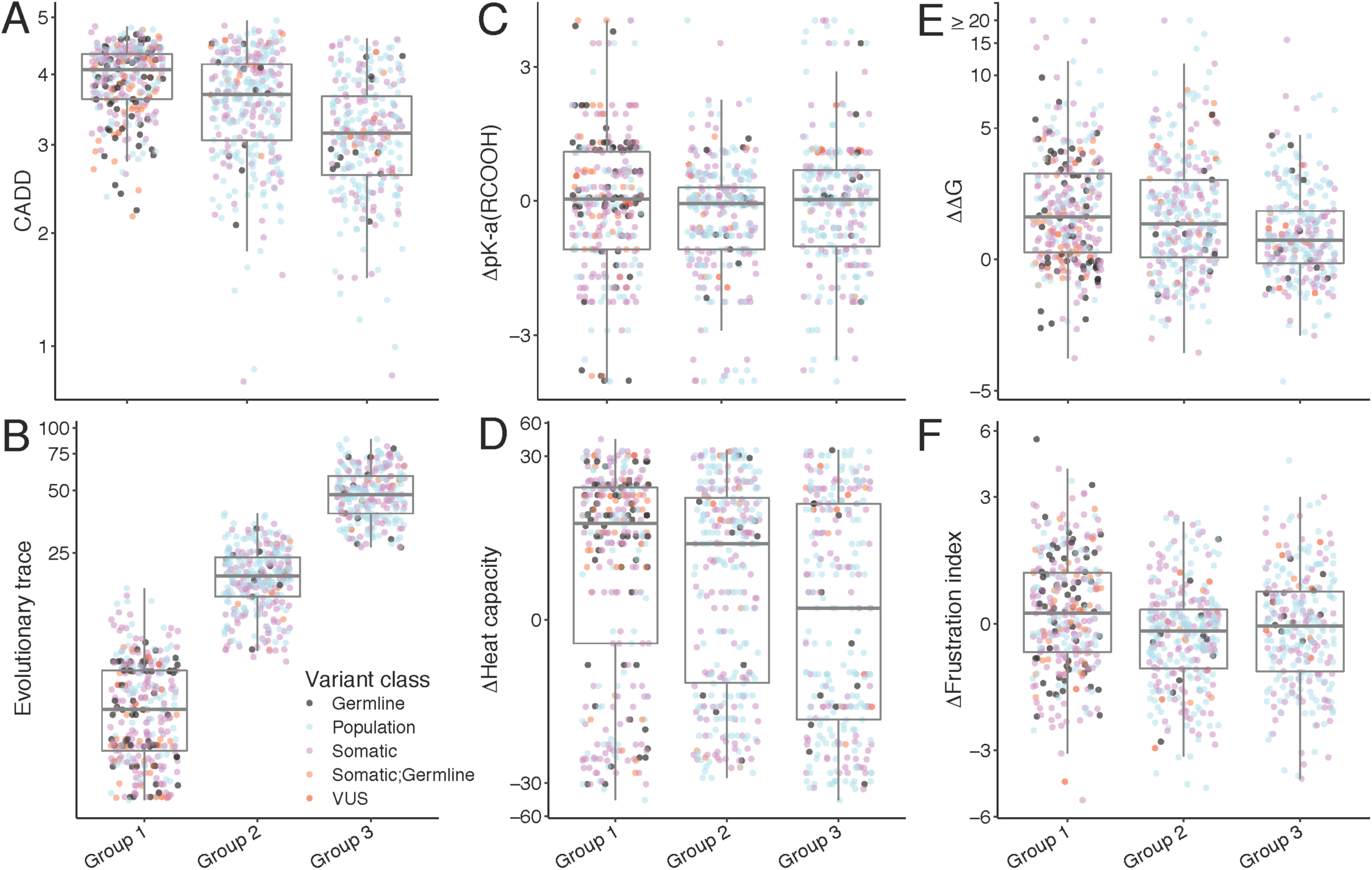
Patterns of scores in the t-SNE topologic groups. **A-F)** Distributions of selected scores shown in box plot for the t-SNE topologic groups for five variant classes (**Figure 5A**). (**A, B**) Sequence-based scores, (**C, D**) amino-acid properties-based scores and (**E, F**) protein structure-based scores.

**Figure S8:**
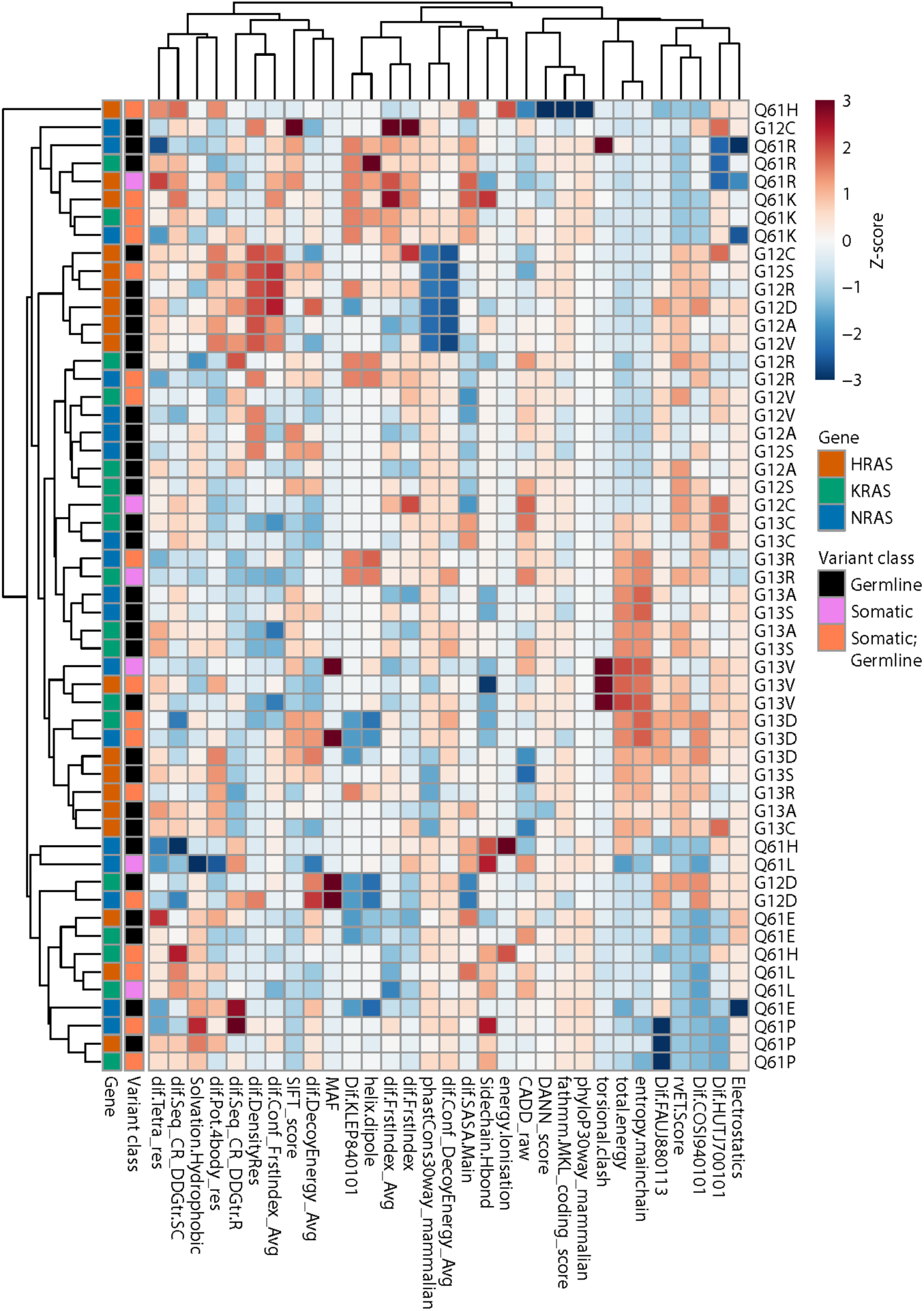
Heatmap plot shows patterns of scores across t-SNE topologic groups for the somatic hotspot variants. Heatmap plot of selected scores are shown in **Figure 6D**.

**Figure S9:**
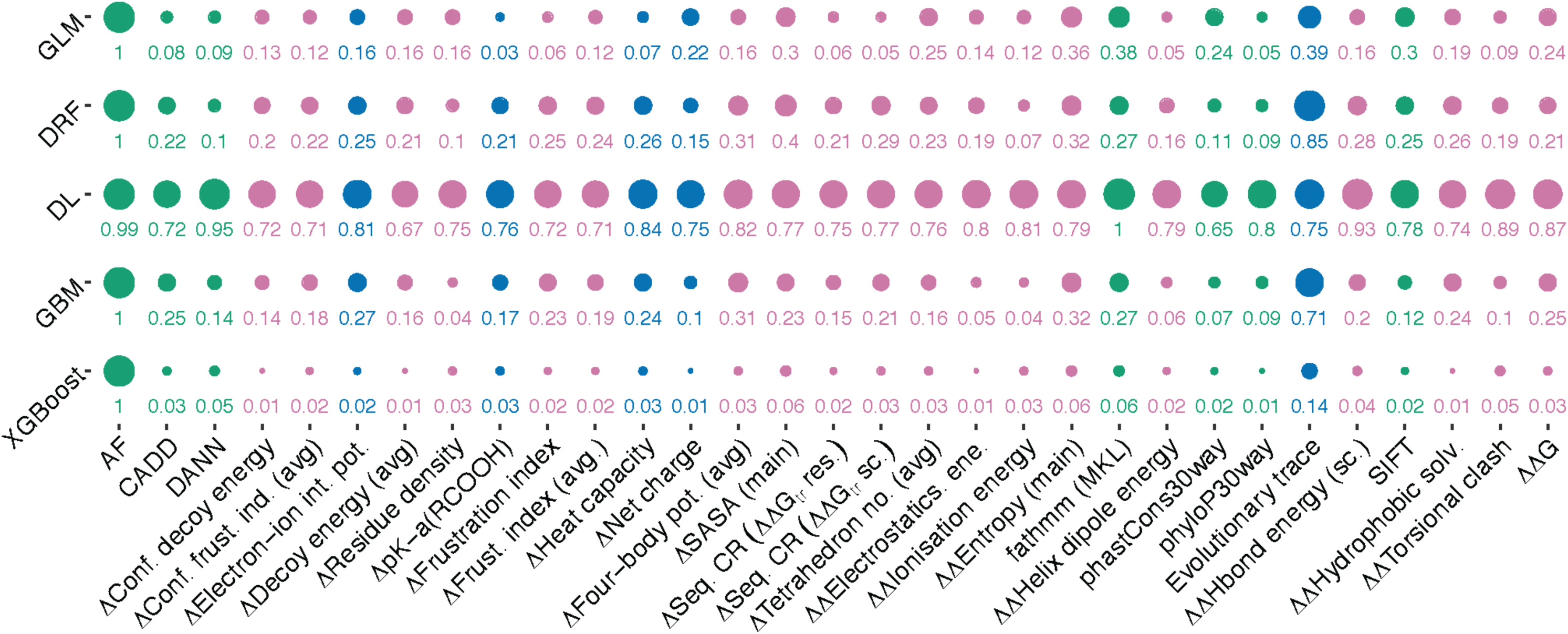
Feature importance from Machine Learning models from the “Best of Family” stacked ensemble. Features are colored by different molecular levels as in **Figure 1**.

**Figure S10:**
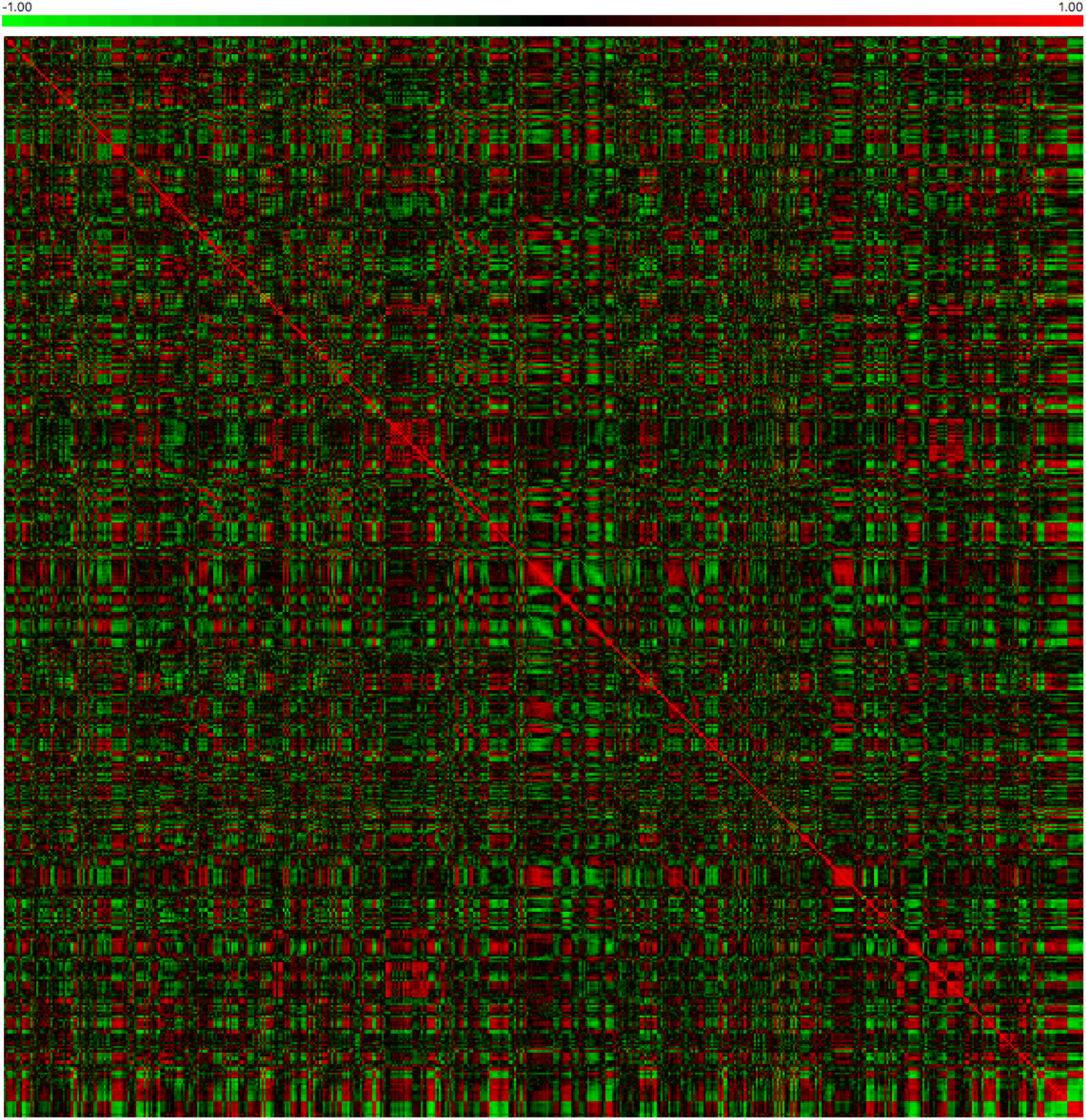
Relationships among amino acid properties of RAS mutations. Spearman correlation of the difference in the 531 AAindex properties between the variants and the WT amino-acid from the Ras sub-family proteins. Each row (and column) is a different variant. The data has been clustered using hierarchical clustering. Color indicates degree of correlation with positive correlations in red and negative in green.

